# Metabolic imprints in the hydrogen isotopes of *Archaeoglobus fulgidus* tetraether lipids

**DOI:** 10.1101/2023.11.29.569324

**Authors:** Jeemin H. Rhim, Sebastian Kopf, Jamie McFarlin, Ashley E. Maloney, Harpreet Batther, Carolynn M. Harris, Alice Zhou, Xiahong Feng, Yuki Weber, Shelley Hoeft-McCann, Ann Pearson, William D. Leavitt

## Abstract

The stable hydrogen isotope composition of archaeal lipids is emerging as a potential paleoenvironmental proxy, adding to the well-established application of plant leaf wax-derived *n*-alkanes in paleohydrological reconstruction. A handful of studies reported relatively invariant and depleted hydrogen isotope compositions for archaeal lipids despite the range of different organisms and growth conditions explored. However, how modes of metabolism and physiological state (growth phase) affect the hydrogen isotope signatures of archaeal lipids remains poorly understood, limiting our ability to interpret archaeal lipid biomarker records from the environment. Here we conducted water isotope label experiments with a metabolically flexible and well-studied model archaeon *Archaeoglobus fulgidus* and quantified the hydrogen isotope fractionation between lipids and water in response to different carbon substrates and electron donor-acceptor pairs. The ^2^H/^1^H fractionation between lipids and water (ε_L/W_) was overall negative, ranging from –280 to –226 ‰, and overlapped with the ranges observed for other archaea in previous studies. Isotope flux-balance model results suggest that ≥80 % and ≥50 % of lipid-bound H in *A. fulgidus* cultures directly reflect water isotope compositions (i.e., not via organic substrate or H_2_) during autotrophy and heterotrophy, respectively. The model results also suggest the final saturation during isoprenoid lipid biosynthesis catalyzed by a flavoenzyme geranylgeranyl reductase likely contributes to the large ^2^H/^1^H fractionation observed in this study, consistent with previous isotope flux-balance model results for a different archaeon. Finally, we synthesized available data to compare ε_L/W_ patterns across all three domains of life: Bacteria, Archaea and Eukarya. Emerging patterns between archaeal and eukaryotic lipids are consistent with the notion of highly fractionating geranylgeranyl reductase, and the patterns between archaeal and bacterial lipids suggest that the general state of energy limitation may also contribute to large, negative values of ε_L/W_ observed in prokaryotic lipids. Altogether, these findings lend further support for the potential of archaeal lipid ε_L/W_ as a paleohydrological proxy and provide a broader insight into the ^2^H/^1^H fractionation mechanisms potentially shared among prokaryotic and eukaryotic lipid biomarkers.

## 1. Introduction

Hydrogen (H) plays a crucial role in most metabolic processes and is incorporated into biological molecules, often with distinct isotope fractionation patterns. Consequently, the ratio of stable hydrogen isotopes (^2^H/^1^H) of certain biomolecules can be used to track the H isotope compositions of source water, or the operation of specific biosynthetic pathways (Leavitt et al., 2023). Lipid-bound H in hydrocarbons may resist isotope exchange over geologic time scales for up to 10^4^ to 10^8^ years (Sessions et al., 2004). Thus, sedimentary records of lipid δ^2^H can be applied to reconstruct past ecological or hydrological states (Sachse et al., 2012; Sessions, 2016; McFarlin et al., 2019). The δ^2^H of lipids produced by photoautotrophic eukaryotes as well as chemoautotrophic and heterotrophic bacteria have been studied in detail for this reason (e.g., Sessions et al., 1999; Chikaraishi and Naraoka, 2003; Zhang and Sachs, 2007; Zhang et al., 2009; Sachse et al., 2012; Sachs, 2014; Dawson et al., 2015; Osburn et al., 2016; Leavitt et al., 2016; Wijker et al., 2019; McFarlin et al., 2019). In contrast, there are only a handful of studies on the H isotope compositions of archaeal lipids (Kaneko et al., 2011; Dirghangi and Pagani, 2013; Wu et al., 2020; Lengger et al., 2021; Leavitt, Kopf et al., 2023). As a consequence, the factors controlling the lipid/water fractionation (ε_L/W_), as well as the extent and limits of proxy applications remain poorly understood for archaeal lipids.

The ε_L/W_ capture a combination of source water δ^2^H and the kinetic isotope effect (KIE) of enzymes involved in lipid biosynthesis. For lipids produced by photoautotrophic eukaryotes (e.g., plants and algae), KIE is relatively invariant but can be altered by some environmental factors (e.g., temperature, salinity, irradiance) and some factors intrinsic to an organism (e.g. growth rate in response to different stimuli and underlying genetics) (Sachs, 2014; Van Der Meer et al., 2015; Sachs and Kawka, 2015; Sachs et al., 2016, 2017; Maloney et al., 2016; Bender, 2017; Ladd and Sachs, 2017; Wolfshorndl et al., 2019). In general, these are well studied in eukaryotes, and have been accounted for in hydroclimate proxy studies (Sachse et al., 2012; Sessions, 2016; McFarlin et al., 2019).

Bacterial lipids show a significantly wider range in ε_L/W_ than eukaryotic lipids. Depending on the central metabolism a given bacteria operates, ε_L/W_ can range from –400 to –200 ‰ (exclusively negative values) for (an)aerobic chemoautotrophs to –154 to +380 ‰, associated with aerobic heterotrophs (Sessions et al., 2002; Campbell et al., 2009; Zhang et al., 2009; Dawson et al., 2015; Osburn et al., 2016; Leavitt et al., 2016; Wijker et al., 2019). For lipids produced by aerobic heterotrophs, H fluxes through NADP^+^-reducing and NADPH-balancing reactions provide a quantitative explanation for the full range of ε_L/W_ values observed (Wijker et al., 2019). Bacterial lipids found in sediments, especially those produced by aerobic heterotrophs, may have more useful applications as proxies for microbial metabolism and ecology rather than past hydroclimate (Zhang et al., 2009; Wijker et al., 2019).

Less is known about what information archaeal ε_L/W_ record. Studies to-date indicate that archaeal lipids are generally depleted in ^2^H, with ε_L/W_ values similar to those observed in eukaryotic photoautotrophs and bacterial chemoautotrophs (Kaneko et al., 2011; Dirghangi and Pagani, 2013; Wu et al., 2020; Lengger et al., 2021; Leavitt, Kopf et al., 2023). In one recent study, continuous cultures of the obligate autotroph *Nitrosopumilus maritimus* revealed that ε_L/W_ remains nearly constant across a 3-fold range in growth rate (Leavitt, Kopf et al., 2023). Other studies have reported similarly large, negative values of ε_L/W_ for archaea grown under heterotrophic conditions (Kaneko et al., 2011; Dirghangi and Pagani, 2013; Wu et al., 2020; Lengger et al., 2021). However, it remains to be determined whether and how carbon metabolism affects ε_L/W_ within a single organism. Testing this is important, considering that mixotrophy (using both autotrophic and heterotrophic carbon sources) is a widespread mode of metabolism in nature and that many archaea are adapted to fluctuating environments. For bacterial lipids, changes in carbon metabolism resulted in significantly different ε_L/W_ values within a single organism for some species (e.g., *Cupriavidus oxalaticus* and *Paracoccus denitrificans*; Zhang et al., 2009; Osburn et al., 2016), but not in other species (sulfate-reducing bacteria; Dawson et al., 2015; Osburn et al., 2016; Leavitt et al., 2016). As such, understanding the role of carbon metabolism in setting archaeal lipid H isotope compositions is important for interpreting archaeal lipid records from the environment. Furthermore, our current understanding of broader patterns of ε_L/W_ that emerge from comparison across lipids produced by all three domains of life—Archaea, Bacteria, and Eukarya—is limited, despite the similarities observed among certain subgroups of different domains.

In this study, we conducted pure culture experiments to better understand factors influencing ε_L/W_ in archaeal lipids. To test the effect of metabolism on ε_L/W_ within a single organism, we cultivate a metabolically flexible archaeon *Archaeoglobus fulgidus* in batch cultures on different carbon substrates and electron donor-acceptor pairs (lactate-thiosulfate, lactate-sulfate, and H_2_/CO_2_-thiosulfate). We also probe the effect of growth phase by comparing biomass harvested at exponential and stationary growth phases and track the fraction of water incorporated into the final product lipid by using different water isotope compositions of growth media. We then interpret the regression parameters derived from water isotope label experiments with an isotope flux-balance model informed by the enzymatic steps involved in the Wood-Ljungdahl (reductive acetyl-CoA) pathway used by *A. fulgidus* as well as archaeal isoprenoid lipid biosynthesis pathways used by all tetraether-producing archaea. In addition to *A. fulgidus*, we examine two thermoacidophilic archaea (*Acidianus* sp. DS80 and *Metallosphaera sedula* DSM 5348T) for additional ε_L/W_ measurements. We synthesize our new findings in the domain Archaea along with other available archaeal lipid ε_L/W_ data as well as representative ε_L/W_ values from domains Bacteria and Eukarya, and highlight factors that underlie patterns in ε_L/W_ across the three domains of life.

## 2. Materials and Methods

### 2.1. Culture Strains and Growth

*Archaeoglobus fulgidus* str. VC-16 (DSM4304) was obtained from Deutsche Sammlung von Mikroorganismen und Zellkulturen (Braunschweig, Germany) and grown in batch cultures on two different carbon substrates and electron acceptors (Table 1). Culture media was prepared following the DSMZ 399 recipe with modifications as follows (per liter): 0.34 g KCl, 4.00 g MgCl_2_×6 H_2_O, 3.45 g MgSO_4_×7 H_2_O, 0.25 g NH_4_Cl, 0.14 g CaCl_2_×2 H_2_O, 0.11 g K_2_HPO_4_×3 H_2_O, 0.20 g KH_2_PO_4_, 18.00 g NaCl, 2.52 g NaHCO_3_, 2 mL Fe(NH_4_)_2_(SO_4_)_2_×7 H_2_O (0.1% w/v), and 10 mL trace elements (DSMZ 141). After autoclaving, sterile anoxic solutions were added to the medium to a final concentration of 3.2 mM L-cysteine-HCl and 1% (v/v) vitamin solution (DSMZ 141). When thiosulfate was used as an electron acceptor, 4.00 g/L MgCl_2_×6 H_2_O and 3.45 g/L MgSO_4_×7 H_2_O were replaced with 4.70 g/L MgCl_2_×6 H_2_O and 3.48 g/L Na_2_S_2_O_3_×5 H_2_O. For heterotrophic growth, lactate was used as an electron donor and carbon source, 20 mM of Na-lactate solution was added and incubated with approximately 1 bar of N_2_/CO_2_ (80:20, v/v) headspace; for autotrophic growth, lactate was omitted and H_2_ and CO_2_ were used as an electron donor and carbon source, respectively, with 21 psig H_2_/CO_2_ (80:20, v/v) in the headspace. The headspaces of autotrophic cultures were flushed and re-pressurized with H_2_/CO_2_ once per day after each OD measurement. The δ^2^H of culture media was manipulated by volumetrically diluting 99.9 % purity ^2^H_2_O.

**Table 1.** Summary of culture experiments.

| Experimental Condition | electron donor | electron acceptor | carbon source |
| --- | --- | --- | --- |
| Thiosulfate-Lactate (T-L) | Lactate (20 mM) | Thiosulfate (14 mM) | Lactate (20 mM) |
| Sulfate-Lactate (S-L) | Lactate (20 mM) | Sulfate (14 mM) | Lactate (20 mM) |
| Thiosulfate-H<sub>2</sub>/CO<sub>2</sub> (T-H<sub>2</sub>/CO<sub>2</sub>) | H<sub>2</sub> (g)* (80% v/v) | Thiosulfate (14 mM) | CO<sub>2</sub> (g)* (20% v/v) |

\* The headspace of autotrophic cultures was filled with 21 psig H<sub>2</sub>/CO<sub>2</sub> (80:20, v/v) and re-pressurized with H<sub>2</sub>/CO<sub>2</sub> once per day after each OD measurement.

All cultures of *A. fulgidus* were incubated at 82 °C. Heterotrophic cultures were incubated statically, and autotrophic cultures were shaken at 200 rpm to increase gas exchange between the H_2_/CO_2_ headspace and culture media. The optical density at 600 nm (OD_600_) of samples was used for monitoring growth, and linearity against direct cell counts was confirmed (depth 0.02 mm; Petroff-Hausser Counter 3900) throughout the absorbance range. Biomass samples were harvested twice per experiment, one during mid-exponential phase and the other during early stationary phase. To harvest, 50 mL of culture was centrifuged for 15 minutes at 4 °C and 4500 *g* and resulting wet cell pellets were stored at –80 °C until lipid extraction.

Batch cultures of *Acidianus* sp. DS80 and *M. sedula* DSM 5348T were grown for additional archaeal lipid ε_L/W_ measurements (not for water isotope labeling experiments or isotope flux-balance model). *Acidianus* sp. DS80 was grown as previously described in (Rhim, Zhou et al., 2024), and *M. sedula* was grown in Brock basal salts medium with Allen’s trace element mixture (Brock et al., 1972). See Supporting Information for additional details for growth conditions for *Acidianus* sp. DS80 and *M. sedula*.

### 2.2. Lipid Extraction, Ether Cleavage, and Quantification

Frozen cell pellets were lyophilized overnight, and dry cell pellets were extracted using acidic hydrolysis-methanolysis. Briefly, dry cell pellets were physically disrupted in 1.5-mL microcentrifuge tubes by vortexing with methanol (MeOH) and *ca*. 250 µL of 100 µm combusted glass beads for 10 minutes at 3000 rpm using a Disruptor Genie (Scientific 126 Industries, SI-DD38). After excess MeOH wasevaporated, octacosanol (C_28_) or lignocerol (C_24_) and C_46_ GTGT were added to all samples as internal extraction standards. Samples were incubated at 65 °C for 90 minutes with 3 N methanolic HCl (33 % water content) to hydrolyze tetraether headgroups. Methyl tert-butyl ether (MTBE) was added to the cooled samples at a ratio of 3:2 (acid:MTBE, v/v), and samples were sonicated for 5 minutes (Qsonica Q500, Newtown, CT, USA). After sonication, n-hexane was added at a 1:1 ratio (MTBE:hexane, v/v), vortexed, and centrifuged at 15,000 g for 3 minutes. The upper organic phase was collected three times with n-hexane for the two subsequent rounds. The resulting total lipid extracts (TLEs) were dried under N_2_ and proceeded to ether cleavage or stored at –20 °C until ether cleavage.

Ether bonds were cleaved in 57% hydroiodic acid (HI) at 125 °C for 4 hours. Then the organic phase was collected three times using n-hexane. The resulting alkyl iodides were reduced to alkanes (biphytanes, BPs) with H_2_ in the presence of Pt^(IV)^O_2_ (Kaneko et al., 2011). Hydrogenated samples were passed through combusted pasteur pipettes with glass wool plugs to remove PtO_2_ powder. Filtered samples were dried under N_2_ and stored at –20 °C until analysis. The resulting ether-cleaved BPs were analyzed on a single quadrupole gas chromatography-mass spectrometer (GC-MS; Thermo ISQ LT with TRACE 1310) for compound identification and on a GC-flame ionization detector (GC-FID; Thermo TRACE 1310) for quantification, both housed in the Organic Geochemistry Laboratory at the University of Colorado Boulder.

### 2.3. Hydrogen isotope analysis and data processing

The δ^2^H of BPs was analyzed on a GC-pyrolysis-isotope ratio MS (GC-P-IRMS) on a GC IsoLink II IRMS System (Thermo Scientific) in the Organic Geochemistry Laboratory at the University of Colorado Boulder. The GC-P-IRMS system consists of a Trace 1310 GC fitted with a programmable temperature vaporization (PTV) injector and either a 30 m ZB5HT column (i.d. = 0.25 mm, 0.25 μm, Phenomenex, Torrance, CA, USA) or a 60 m DB1 column (i.d. = 0.25 mm, 0.25 μm, Agilent, Santa Clara, CA, USA), ConFlo IV interface, and MAT 253 Plus mass spectrometer (Thermo Scientific).

*A. fulgidus* has been observed to produce both tetraether and diether lipids, with the majority of GDGT with no ring (GDGT-0) and a small fraction with up to two pentacyclic rings (Lai et al., 2008). The tetraether-to-diether ratio for *A. fulgidus* is expected to be ∼1:1 at the growth temperature in this study (82 °C; Lai et al., 2008), and we focus on the tetraether fraction for the rest of the discussion. In line with the previous observations, the majority of BPs derived from *A. fulgidus* GDGT did not have rings (BP-0). All δ^2^H values reported in this study are for BP-0.

All hydrogen isotope ratios are reported in the standard delta notation (δ^2^H) in permil (‰) units, against the Vienna Standard Mean Ocean Water (VSMOW) international standard:

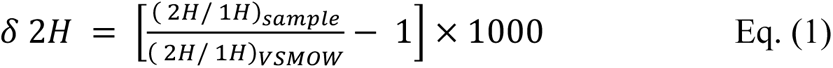

The lipid-water hydrogen isotope fractionation factor is reported in alpha notation (^2^α_L/W_), and fractionation is reported in epsilon notation (^2^ε_L/W_) in ‰:

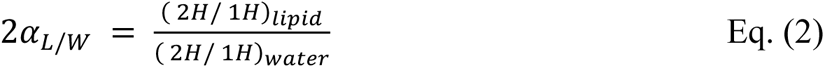

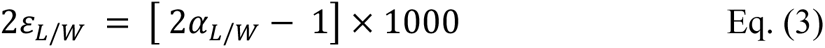

Values of δ^2^H were first determined relative to H_2_ reference gas (δ^2^H_raw_), and then calibrated externally using a standard *n*-alkane mixture (A6, containing C_15_ through C_30_ n-alkanes spanning –9 to –263 ‰ vs. VSMOW; A. Schimmelmann, Indiana University). The A6 standard was combined with a C_36_ n-alkane (nC_36_, –259.2 ‰ vs. VSMOW; A. Schimmelmann, Indiana University) and measured at regular intervals at different concentrations. The BP δ^2^H calibration was done in R, using the packages *isoreader* (Kopf et al., 2021) and *isoprocessor* available at github.com/isoverse. Details of the data reduction and calibration methods are described in (Leavitt, Kopf et al., 2023). The RMSE of the calibration peaks was 4.2 ‰ (n=613) across a peak size range from 6.2 to 59.9 Vs. The RMSE of the C_36_ *n*-alkane internal standard was 7.5 ‰ (n=90) across a peak size range from 6.4 to 50.9 Vs. Calibrated δ^2^H values for the biphytanes were corrected for the H added during hydrogenation of alkyl iodides (see Leavitt, Kopf et al., 2023). The hydrogenation correction ranged from 9.8 to 13.7 ‰ and increased analytical uncertainty by up to 1.8 ‰.

The δ^2^H of the H_2_ gas in the H_2_/CO_2_ headspace of autotrophic cultures was analyzed on a Thermo Gasbench II in the Stable Isotope Laboratory at Northwestern University. The δ^2^H of non-exchangeable H in lactate used for heterotrophic cultures was assumed to be –60 ‰, which is the value previously determined for glucose (Zhang et al., 2009).

The δ^2^H of natural abundance and deuterium-enriched medium water was measured at the Stable Isotope Laboratory at Dartmouth College as previously described (Kopec et al., 2019). Briefly, water was reduced with chromium at 850 °C using the H-Device, and the δ^2^H of the resulting gas was measured by an IRMS (Thermo Delta Plus XL). The measured δ^2^H values were converted to the water δ^2^H equivalent by calibrating with water standards with known isotopic compositions analyzed in the same way. For samples expected to be outside the range of the standards (e.g., the deuterium-enriched medium), samples were diluted with water standards. The 1σ uncertainty for δ^2^H measurements is <0.5 ‰.

### 2.4. Model implementation

An isotope flux-balance model was adapted from the conceptual framework described in Leavitt, Kopf et al. (2023). Our model considers the Wood-Ljungdahl (reductive acetyl-CoA) pathway used by A. fulgidus for determining the sources and fluxes of H during acetyl-CoA (Ac-CoA) synthesis. It also considers the general archaeal lipid biosynthesis pathway for the accounting of total H budget in archaeal lipids. Analytical solutions for the model were calculated in R. Details of the model design and outcomes are described in section 4.

## 3. Results

### 3.1. Cultures and growth rates

*Archaeoglobus fulgidus* is a metabolically flexible strain from the phylum Euryarchaeota. We cultivated *A. fulgidus* in batch (closed-system) cultures on three different combinations of electron donors and acceptors (Table 1; Fig. 1A–C). In two treatments, *A. fulgidus* was grown as a heterotroph, using lactate as the carbon source and electron donor and sulfate or thiosulfate as the terminal electron acceptor. In the third treatment, *A. fulgidus* was grown as an autotroph, using CO_2_ as the carbon source, H_2_ as the electron donor, and thiosulfate as the terminal electron acceptor. Cultures were acclimated to each treatment by growing on respective substrate combinations for two or more passages before inoculation. Overall, heterotrophic conditions resulted in faster growth (T-L, *t*_d_ = 6.2 ± 0.6 h; S-L, *t*_d_ = 10.3 ± 1.7 h) compared to autotrophic growth (T-H_2_/CO_2_, *t*_d_ = 19.3 ± 3.9 h) (Table 2; Fig. 1A–C).

**Figure 1.**
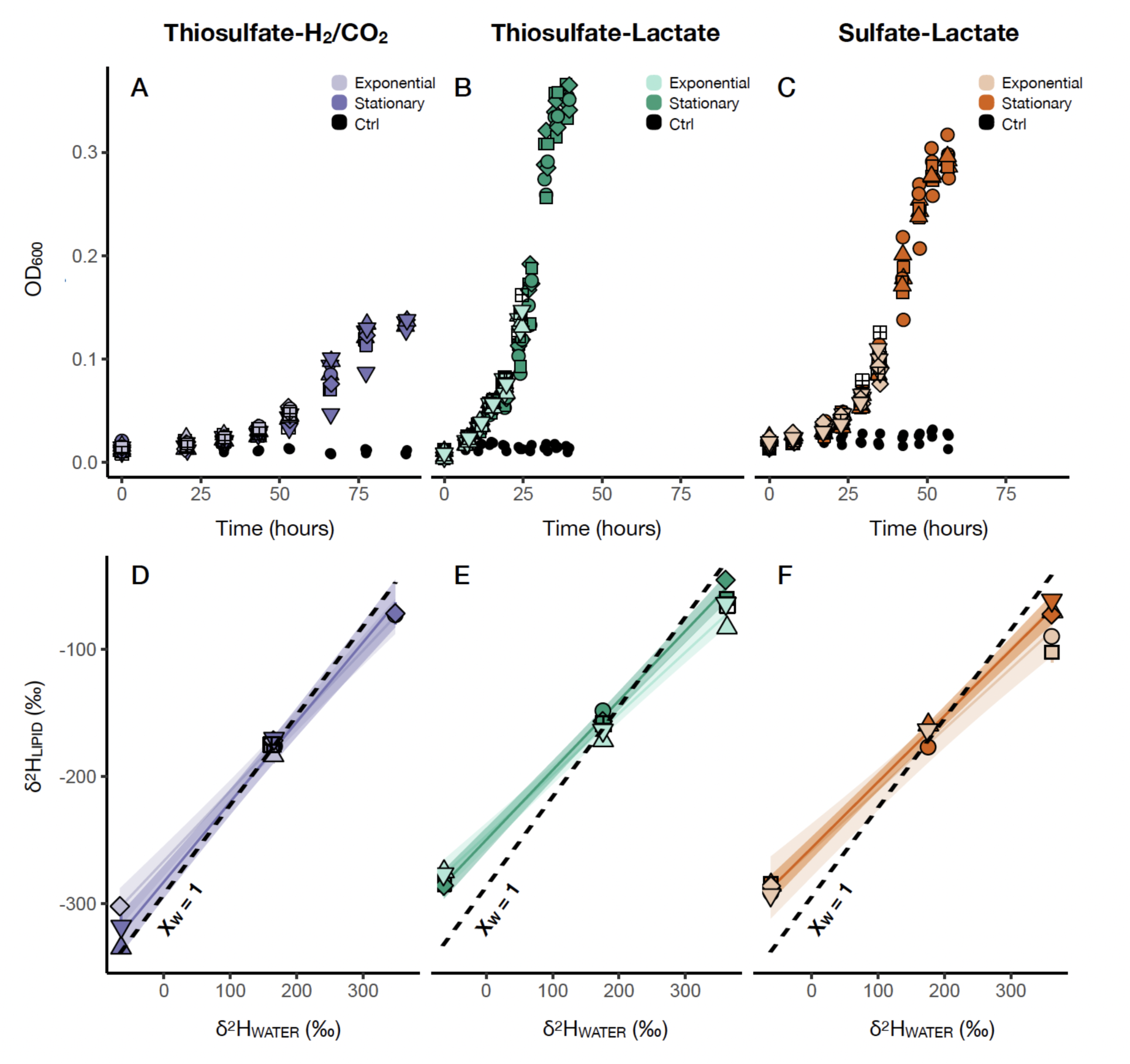
Growth curves and regression of lipid δ^2^H versus water δ^2^H for *Archaeoglobus fulgidus*. Growth curves of *A. fulgidus* grown autotrophically on (A) thiosulfate and H_2_/CO_2_ or heterotrophically on (B) thiosulfate and lactate or (C) sulfate and lactate. Regression of biphytane hydrogen isotopic composition (δ^2^H_L_) on water isotopic composition (δ^2^H_W_) grown on (D) thiosulfate and H_2_/CO_2_, (E) thiosulfate and lactate, and (F) sulfate and lactate.

**Table 2.** Growth and isotope data for tetraether-derived biphytanes produced by *Archaeoglobus fulgidus* grown on different carbon substrates and electron donor-acceptor pairs.

| Substrate<sup>a</sup> | $t_d$ (h)<sup>b</sup> | Growth Phase<sup>c</sup> | Avg. OD<sup>d</sup> | $\delta^2H_W^e$ (‰) | s.d. (‰) | $\delta^2H_L^f$ (‰) | s.d. (‰) | $^2\epsilon_{L/W}$ (‰) | s.d. (‰) | $n^g$ | Slope<sup>h</sup> ($^2\alpha_{L/W} \cdot f_W$) | s.e. | R<sup>2</sup> | $p$ |
| --- | --- | --- | --- | --- | --- | --- | --- | --- | --- | --- | --- | --- | --- | --- |
| T-L | $6.2 \pm 0.6$ | Exp | 0.147 | 362.4 | 0.2 | -70.7 | 6.6 | -317.9 | 29.4 | 3 | 0.481 | 0.017 | 0.996 | $1 \times 10^{-7}$ |
|  |  |  | 0.135 | 176.0 | 0.2 | -164.6 | 7.0 | -289.6 | 12.4 | 3 |  |  |  |  |
|  |  |  | 0.141 | -64.0 | 0.2 | -275.8 | 7.1 | -226.2 | 5.9 | 2 |  |  |  |  |
| | | Sta | 0.346 | 361.3 | 0.2 | -52.7 | 6.7 | -304.1 | 38.5 | 2 | 0.548 | 0.015 | 0.999 | $3 \times 10^{-6}$ |
|  |  |  | 0.344 | 175.5 | 0.2 | -152.4 | 7.5 | -279.0 | 13.7 | 2 |  |  |  |  |
|  |  |  | 0.351 | -63.7 | 0.2 | -285.3 | 8.7 | -236.7 | 7.2 | 2 |  |  |  |  |
| S-L | $10.3 \pm 1.7$ | Exp | 0.096 | 361.8 | 0.2 | -87.5 | 9.7 | -329.9 | 36.4 | 1 | 0.477 | 0.028 | 0.993 | $7 \times 10^{-5}$ |
|  |  |  | 0.100 | 174.7 | 0.2 | -163.5 | 4.8 | -287.9 | 8.5 | 3 |  |  |  |  |
|  |  |  | 0.082 | -61.7 | 0.2 | -289.7 | 5.9 | -243.0 | 5.0 | 2 |  |  |  |  |
| | | Sta | 0.307 | 361.4 | 0.2 | -66.9 | 5.7 | -314.6 | 26.9 | 2 | 0.519 | 0.016 | 0.998 | $4 \times 10^{-7}$ |
|  |  |  | 0.295 | 175.4 | 0.2 | -168.1 | 4.9 | -292.3 | 8.5 | 2 |  |  |  |  |
|  |  |  | 0.282 | -61.5 | 0.2 | -287.1 | 10.2 | -240.3 | 8.6 | 2 |  |  |  |  |
| T-H<sub>2</sub>/CO<sub>2</sub> | $19.3 \pm 3.9$ | Exp | 0.052 | 348.3 | 0.2 | -72.8 | 5.0 | -312.3 | 21.5 | 2 | 0.552 | 0.019 | 0.998 | $8 \times 10^{-5}$ |
|  |  |  | 0.046 | 164.8 | 0.2 | -176.8 | 7.0 | -294.0 | 11.5 | 1 |  |  |  |  |
|  |  |  | 0.054 | -66.5 | 0.2 | -302.1 | 4.7 | -252.4 | 4.0 | 1 |  |  |  |  |
| | | Sta | 0.137 | 348.5 | 0.2 | -71.7 | 8.7 | -311.7 | 37.8 | 3 | 0.628 | 0.025 | 0.997 | $1 \times 10^{-5}$ |
|  |  |  | 0.132 | 166.0 | 0.2 | -173.8 | 10.0 | -291.4 | 16.8 | 1 |  |  |  |  |
|  |  |  | 0.134 | -64.5 | 0.2 | -326.4 | 10.2 | -279.9 | 8.8 | 2 |  |  |  |  |
| T-H<sub>2</sub>/CO<sub>2</sub> (calculated) | | - | - | - | - | - | - | - | - | - | 0.707 ($f_W = 1$) | - | - | - |

<sup>a</sup> Thiosulfate-lactate (T-L); sulfate-lactate (S-L); thiosulfate-H<sub>2</sub>/CO<sub>2</sub> (T-H<sub>2</sub>/CO<sub>2</sub>)
<sup>b</sup> Doubling time calculated for logarithmic growth
<sup>c</sup> Biomass harvested during mid-exponential growth phase (Exp); biomass harvested during early stationary phase, including biomass from both exponential and stationary phase (Sta)
<sup>d</sup> Optical density (OD)
<sup>e</sup> $\delta^2H$ of culture medium at the time of biomass sampling. Average of biological and technical replicates $\pm 1$ s.d.
<sup>f</sup> Weighted mean $\delta^2H$ of biphytane from GDGT-0
<sup>g</sup> Number of biological replicates used to calculate the average values of $\delta^2H_{Water}$ , $\delta^2H_{Lipid}$ , and $^2\epsilon_{L/W}$
<sup>h</sup> Slope for the regression of $\delta^2H_{Lipid}$ against $\delta^2H_{Water}$ (see Figure 1D-F)

### 3.2. Carbon metabolism and growth phase affect lipid δ^2^H and regression parameters

Different modes of carbon metabolism in *A. fulgidus* not only resulted in different growth rates but also affected lipid δ^2^H and regression parameters. The lipid isotopic composition reflects the sum of distinct isotope fractionations between lipids and each external H source, resulting in the isotopic mass balance (Zhang et al., 2009):

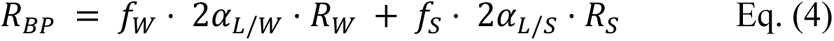

where R_BP_, R_W_, and R_S_ represent the ^2^H/^1^H ratios of BPs from GDGT-0 (BP-0), media water, and substrate (lactate for the heterotrophic conditions; H_2_ for the autotrophic condition), respectively. *f*_W_ and *f*_S_ are the fractions of lipid-bound H originating from water and substrate, respectively. ^2^α represents the net isotope fractionation factor between lipid and water (^2^α_L/W_) and between lipid and substrate (^2^α_L/S_), respectively. In parallel cultures where only one parameter (R_W_) is varied experimentally, measurements of R_L_, R_W_, and R_S_ provide the basis for regression of R_L_ on R_W_. The isotope analysis results and regression parameters are summarized in Table 2.

The δ^2^H_L_ values were strongly correlated with δ^2^H_W_ values in all experiments (Table 2; R^2^ ≥ 0.993 and *p* ≤ 0.05 in every regression). The regression slopes varied as a function of carbon metabolism and growth phase. Overall, the two heterotrophic conditions (T-L and S-L) and exponential growth phase resulted in lower slopes compared to the autotrophic condition (T-H_2_/CO_2_) and stationary growth phase. Slopes for the two heterotrophic conditions were similar (0.481 ± 0.017 for T-L, 0.477 ± 0.028 for S-L during exponential phase; 0.548 ± 0.015 for T-L, 0.519 ± 0.016 for S-L during stationary phase; Table 2). The autotrophic condition resulted in higher slopes (0.552 ± 0.019, exponential phase; 0.628 ± 0.025, stationary phase; Table 2). Notably, the higher slope observed during the stationary phase of the autotrophic culture was the closest to the slope estimated assuming *f*_W_ = 1 (0.707) (Table 2; Fig. 1D).

## 4. *A. fulgidus* H-isotope flux-balance model

To further interpret these findings, we modified the isotope flux-balance model recently developed for an ammonia-oxidizing archaeon *N. maritimus* SCM1 (Leavitt, Kopf et al., 2023) to reflect the biochemistry underlying the different modes of carbon metabolism in *Archaeoglobus fulgidus* VC-16. The overall conceptual framework consists of two parts for the stoichiometric accounting of: 1) methyl-H in Ac-CoA and 2) the 80 H atoms in the C_40_ alkyl chain of BPs inherited from Ac-CoA. The former requires metabolism-specific information considering the mode(s) of carbon fixation and experimental conditions such as substrate availability, while the latter is informed by more universal information about archaeal isoprenoid biosynthesis. Below, we describe the details of the two parts of our model.

### 4.1. Sources of lipid-H via the modified Wood-Ljungdahl pathway

#### 4.1.1. Heterotrophy

*A. fulgidus* can grow autotrophically or heterotrophically via the modified reductive Ac-CoA or Wood-Ljungdahl pathway (Möller-Zinkhan et al., 1989; Möller-Zinkhan and Thauer, 1990; Hocking et al., 2014). During chemoorganoheterotrophic growth on lactate, the type species *A. fulgidus* VC16 couples the complete oxidation of organic substrates to CO_2_ with the reduction of sulfate or thiosulfate to sulfide. In this study, the organic substrate was lactate. *A. fulgidus* oxidizes lactate via pyruvate and Ac-CoA using enzymes and cofactors involved in the C_1_-pathway, similar to those found in methanogens, in reverse direction (Fig. 2A; Möller-Zinkhan et al., 1989; Möller-Zinkhan and Thauer, 1990). A D-lactate dehydrogenase in *A. fulgidus* catalyzes the first step of lactate oxidation to pyruvate coupled with the reduction of menaquinone (MQ), resulting in menaquinol (MQH_2_) (Step 1, Fig. 2A; Kunow et al., 1993; Reed and Hartzell, 1999). The next step of pyruvate oxidation to Ac-CoA is catalyzed by pyruvate:ferredoxin oxidoreductase (Step 2, Fig. 2A; Kunow et al., 1993). The three H atoms in the methyl group of lactate most likely remain intact and are inherited to Ac-CoA. Some of these Ac-CoA products are used for biomass synthesis (Step 3.2, Fig. 2A), including isoprenoid lipid biosynthesis, while some of them are degraded further for the complete oxidation of lactate. For the latter, carbon monoxide dehydrogenase catalyzes both the reversible carbon-carbon cleavage of Ac-CoA and the oxidation of carbonyl group to CO_2_ coupled with ferredoxin (Fd) reduction (Step 3.1, Fig. 2A; Möller-Zinkhan and Thauer, 1990).

**Figure 2.**
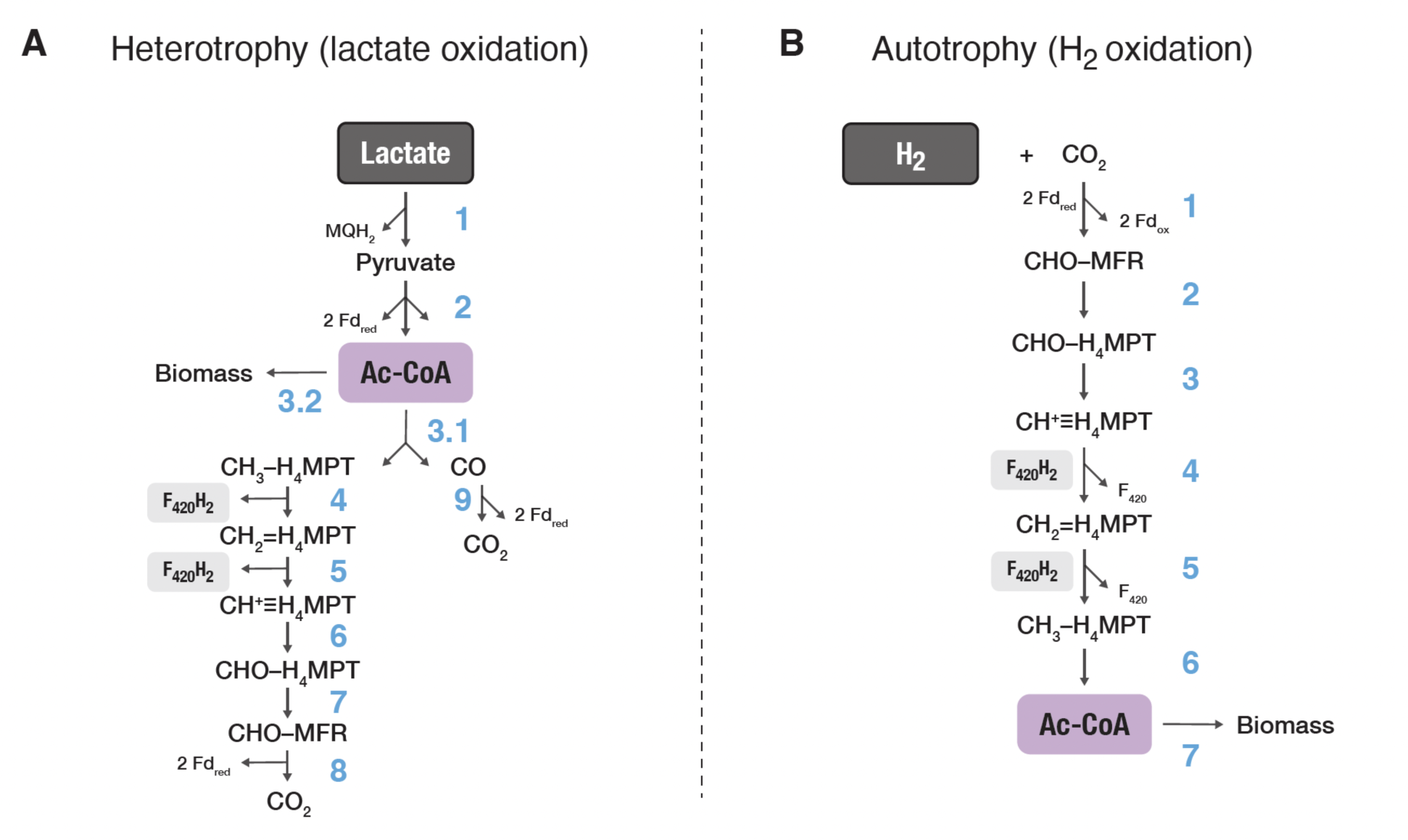
Carbon metabolism by *Archaeoglobus fulgidus* via the modified reductive acetyl-CoA or Wood-Ljungdahl pathway. (A) Chemoorganoheterotrophic growth on lactate: lactate is oxidized via pyruvate and acetyl-CoA, which either serves as the building blocks for biosynthesis or is further degraded for the complete oxidation of lactate. (B) Chemolithoautotrophic growth on H_2_/CO_2_: CO_2_ is reduced to produce acetyl-CoA via the C_1_-pathway, with enzymes and cofactors similar to those involved in methanogenesis. Abbreviations: MQH_2_, menaquinol or reduced menaquinone; Fd_red_, reduced ferredoxin; Ac-CoA, acetyl-CoA; CH_3_-H_4_MPT, methyl-tetrahydromethanopterin (H_4_MPT); CH_2_=H_4_MPT, methylene-H_4_MPT; CH^+^≡H_4_MPT, methenyl-H_4_MPT; CHO-H_4_MPT, formyl-H_4_MPT; CHO-MFR, formylmethanofuran; F_420_H_2_, reduced coenzyme F_420_.

The coenzymes involved in the downstream oxidative branch of Ac-CoA degradation could transfer hydrides (e.g., F_420_H_2_ in Steps 4 and 5, Fig. 2A). These fluxes of H are important to track because the hydrides from Ac-CoA (ultimately from lactate) can be recycled for NADP reduction, the product of which (NADPH) can donate hydrides to isoprenoid precursors during lipid biosynthesis. We assign F_420_ and Fd as the electron acceptors for Steps 4–5 and Step 8, respectively (Fig. 2A), given that the genes relating to all steps of the reductive Ac-CoA pathway are expressed in *A. fulgidus* (Hocking et al., 2014) and considering the known mechanisms of how the enzymes of interest operate in methanogens. In CO_2_-reducing methanogens, both methylenetetrahydromethanopterin reductase (Mer) and methylenetetrahydromethanopterin dehyrogenase (Mtd) oxidize F_420_H_2_ (Steps 4 and 5, respectively, in reverse; Fig. 2A), and formylmethanofuran dehydrogenase (Fwd) requires Fd_red_ (Step 8, in reverse; Fig. 2A) (Thauer, 2012).

#### 4.1.2. Autotrophy

During chemolithoautotrophic growth *A. fulgidus* VC16 couples H_2_ or formate oxidation with thiosulfate or sulfite reduction (Achenbach-Richter et al., 1987; Steinsbu et al., 2010). We fed *A. fulgidus* H_2_ and thiosulfate as the electron donor and acceptor for the autotrophic condition (T-H_2_/CO_2_). *A. fulgidus* reduces CO_2_ to synthesize Ac-CoA via the C_1_-pathway (Fig. 2B), in principle a reversal of the Ac-CoA degradation during heterotrophy (Fig. 2A). The same enzymes involved in H additions used by CO_2_-reducing methanogens catalyze the successive addition of H during autotrophic carbon fixation in *A. fulgidus*: Fwd (Step 1), Mtd (Step 4), and Mer (Step 5) (Fig. 2B). During chemolithoautotrophy, the H atoms in methyl precursors in theory derive from intracellular water. However, one of the isoenzymes for methylenetetrahydromethanopterin (CH_2_-H_4_MPT) dehydrogenase is H_2_-dependent (Hmd) and uses H_2_ directly, as opposed to F_420_H_2_ used by the other isoenzyme Mtd, to reduce methyl-H_4_MPT (Hendrickson and Leigh, 2008). Given the similarities between methanogens and *A. fulgidus*, we consider the possible flux of H from H_2_ to Ac-CoA (Step 4–6, Fig. 2B) in one of the end-member scenarios (see 4.2; Table 3).

**Table 3.**
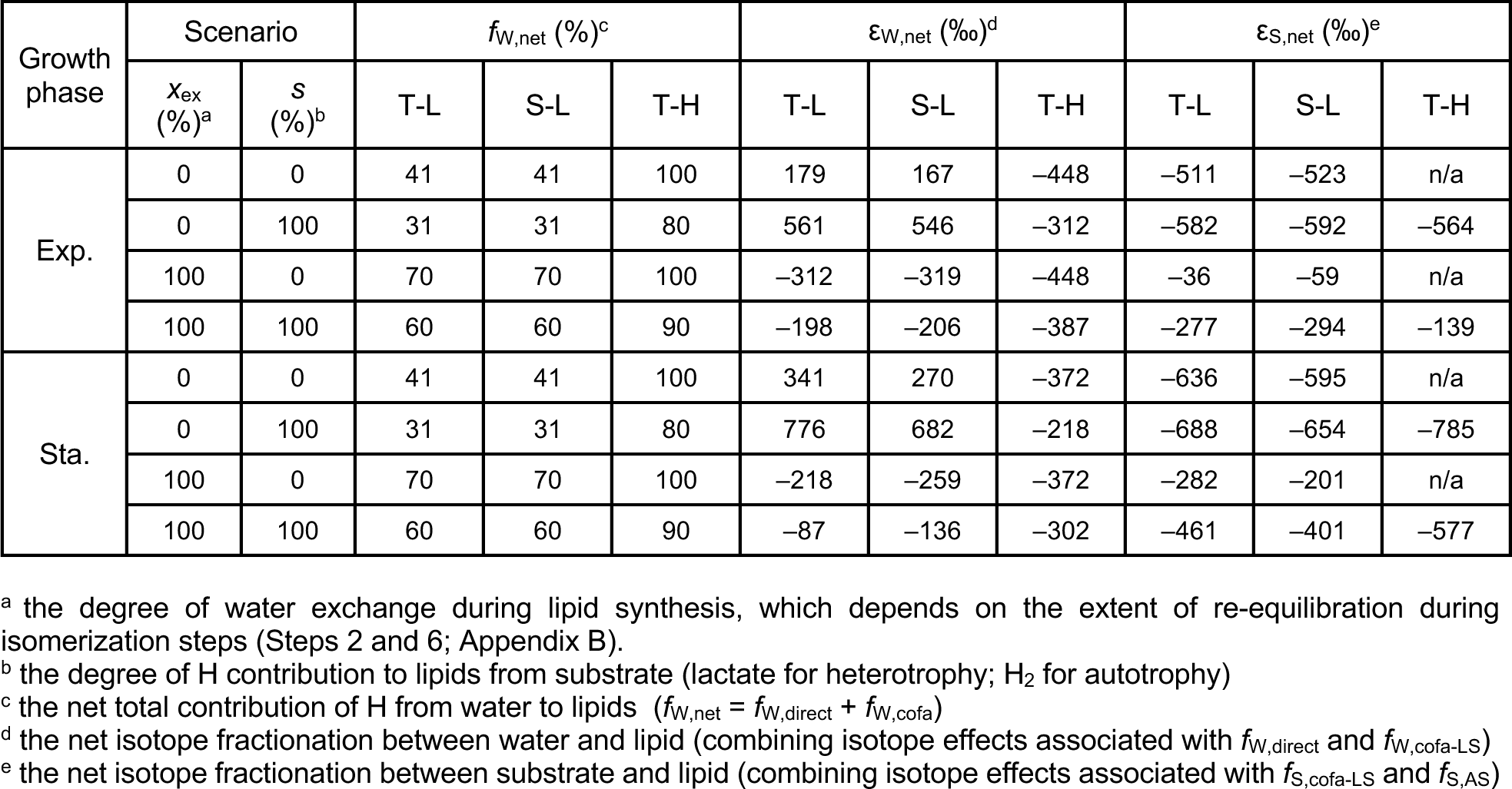
Isotope flux-balance model results for *Archaeoglobus fulgidus*.

### 4.2. Model schematics and stoichiometric accounting

Fractions of Ac-CoA produced via the modified Wood-Ljungdahl pathway described above are used for the synthesis of archaeal isoprenoids. Leavitt, Kopf et al. (2023) provide a detailed overview of the sources of H and mechanisms of the biochemical steps involved in archaeal tetraether biosynthesis (Appendix B). In short, the mevalonate pathway synthesizes isopentenyl pyrophosphate (IPP), which are C_5_ isoprene units, from three units of Ac-CoA. Then, three units of IPP and its isomer condense to yield the C_20_ precursor, geranylgeranyl diphosphate (GGPP). The geranylgeranyl chains are transferred around to yield di-O-geranylgenranylglyceryl phosphate (DGGGP) with the glycerol-1-phosphate (G1P) backbone. The tetraether synthase (Tes) enzyme catalyzes the condensation of two DGGGP units to yield the membrane-spanning tetraether. Finally, the geranylgeranyl reductase (GGR) enzyme catalyzes the saturation of tetraether to yield GDGT. We combine the stoichiometric accounting of the methyl-H in Ac-CoA for *A. fulgidus* (see 4.1) and of the 80 H atoms in the C_40_ alkyl chain of BPs inherited from Ac-CoA during tetraether biosynthesis described above (Leavitt, Kopf et al., 2023) to build an overall schematic of the specific model scenarios explored in this study (Fig. 3).

**Figure 3.**
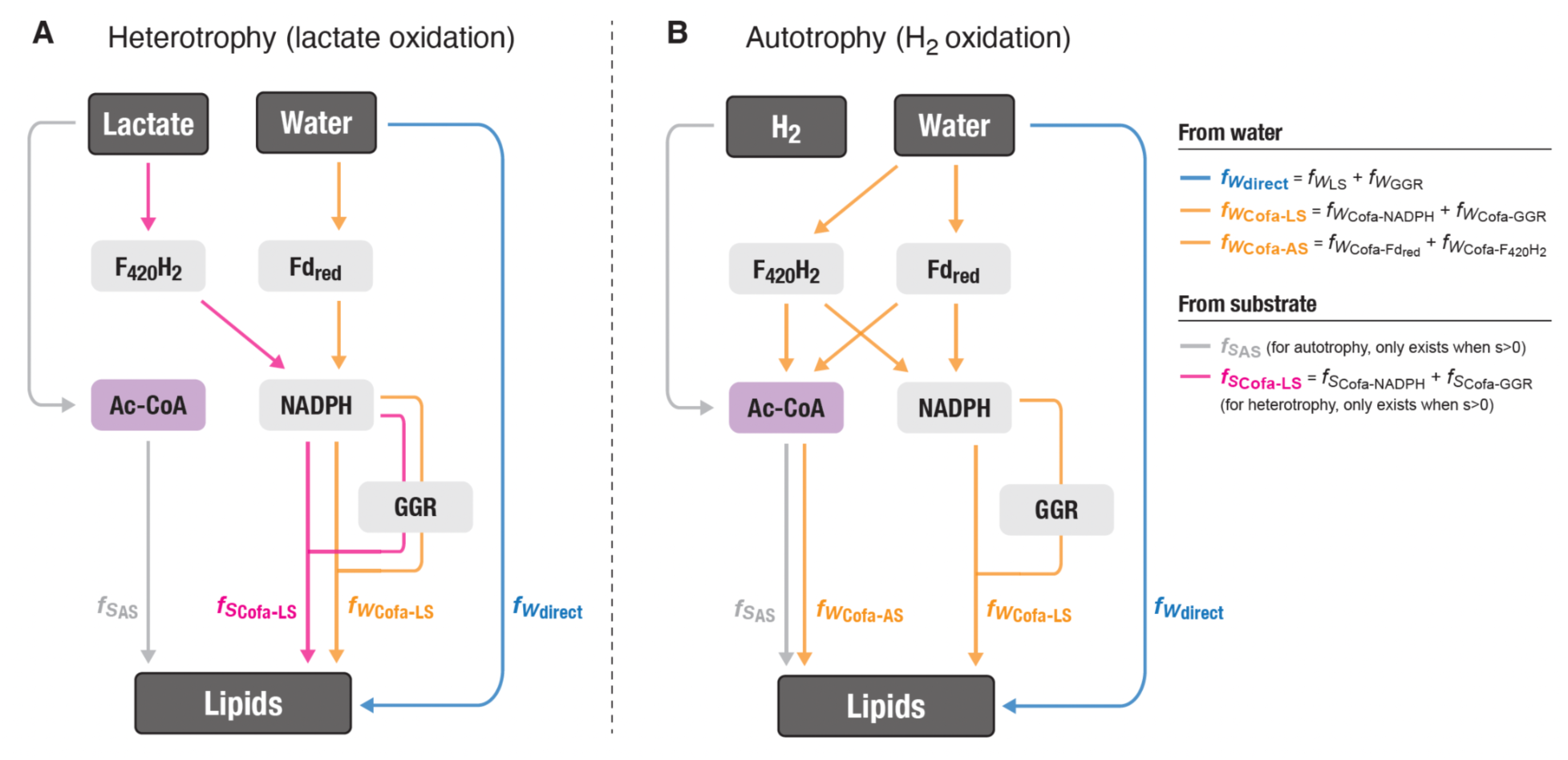
Schematic diagram of the isotope flux-balance model for *Archaeoglobus fulgidus*. (A) Heterotrophic scenario for lactate oxidation; (B) autotrophic scenario for H_2_ oxidation. Color codes for the different H fluxes (*f*) are shown in the legend. The subscript W stands for water as the ultimate H source; the subscript S stands for substrate as the ultimate H source, where the substrates are lactate and H_2_ for heterotrophy and autotrophy, respectively. See 4.2.1 and Appendix A (Eq. A5–A8) for detailed descriptions of individual fluxes. Abbreviations: LS, lipid synthesis; AS, acetyl-CoA synthesis; Cofa, cofactor; F_420_H_2_, reduced coenzyme F_420_; Fd_red_, reduced ferredoxin; Ac-CoA, acetyl-CoA; GGR, geranylgeranyl reductase.

#### 4.2.1. Empirically informed isotopic mass balance

With the fractions defined in the overall hydrogen budget (see Appendix A), the isotopic mass balance in Eq. (4) can be further refined. Replacing the *f*_W_ and *f*_S_ terms in Eq. (4) with the sums of all fractions in Eq. (A5) (Appendix A), isotopic mass balance for the heterotrophic case becomes:

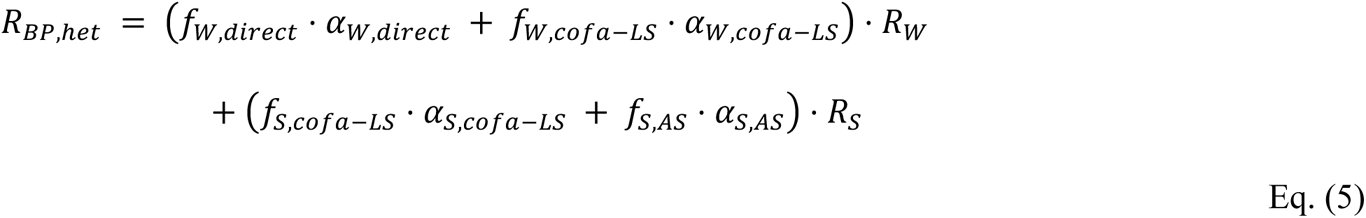

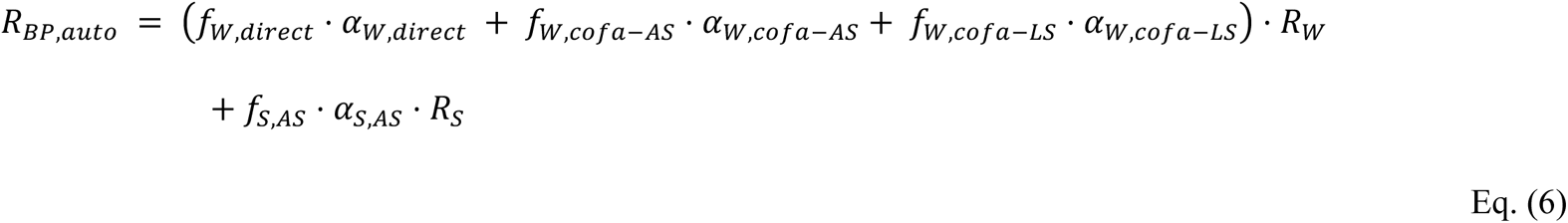

Now we can use the regression parameters estimated from the δ^2^H_H2O_ label experiments (Fig. 1; Table 2) to calculate analytical solutions for reasonable ranges of individual terms in Eq. (5–6). In Eq. (5–6), the coefficient for R_W_ in the first term represents the slope, *m*, and the entire second term represents the intercept, *c*. For heterotrophy:

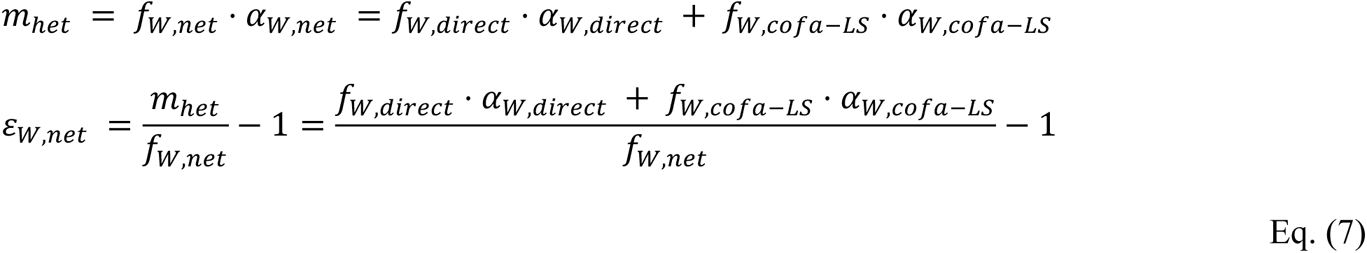

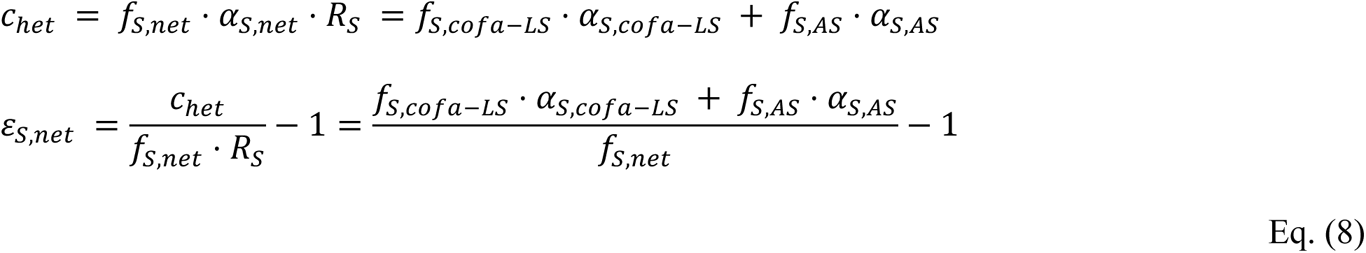

where ε_W,net_ and ε_S,net_ refer to the net isotope fractionation between water and lipid (combining isotope effects associated with *f*_W,direct_ and *f*_W,cofa-LS_) and between substrate and lipid (combining isotope effects associated with *f*_S,cofa-LS_ and *f*_S,AS_), respectively.

### 4.3. Model results

Table 3, Fig. 4 and Fig. 5 show the summary of model results. We first estimated *f*_W,net_, which we define as the total flux that directly or indirectly (e.g., via cofactors) reflects water isotope composition (i.e., not affected by substrate isotopic composition) (*f*_W,net_ = *f*_W,direct_ + *f*_W,cofa_), for a range of *x*_ex_ from 0 % to 100 %. Overall, the autotrophic case had a higher range of *f*_W,net_ compared to the two heterotrophic cases (Table 3; Fig. 4A–C). Even for the minimal water exchange and maximal substrate contribution scenario (*x*_ex_ = 0 %, *s* = 100 %), when the contribution from substrate is expected to be the highest, *f*_W,net_ was 80 % (Table 3; dashed line, Fig. 4A). For the two heterotrophic cases, the *f*_W,net_ values were 31 and 70 % for the highest (*x*_ex_ = 0 %, *s* = 100 %) and lowest (*x*_ex_ = 100 %, *s* = 0 %) substrate contribution scenarios, respectively (Table 3; Fig. 4B–C).

**Figure 4.**
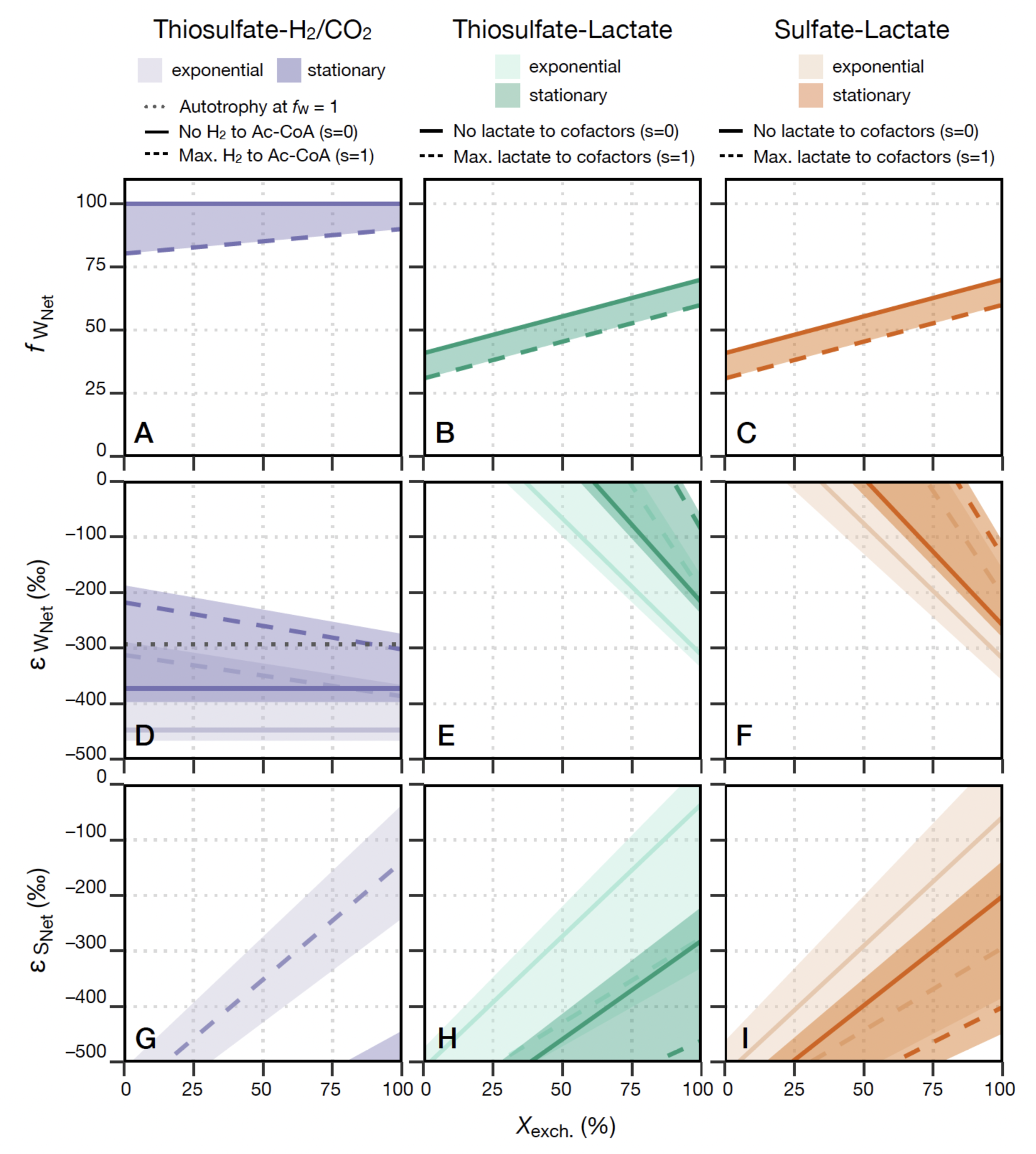
Isotope flux-balance model results for *Archaeoglobus fulgidus*. (A–C) The total flux that directly or indirectly (e.g., via cofactors) reflects water isotope composition (i.e., not affected by substrate isotopic composition) (*f*_W,net_); (D– F) Net isotope fractionation between lipid and water (ε_W,net_); (G–I) Net isotope fractionation between lipid and substrate (ε_S,net_). Each column represents the metabolic mode and growth substrates: autotrophy on thiosulfate and H_2_/CO_2_ (A, D and G); heterotrophy on thiosulfate and lactate (B, E and H); and heterotrophy on sulfate and lactate (C, F and I). Lighter shaded areas refer to exponential growth phase; darker shades refer to stationary growth phase. Solid lines refer to the s = 0 % (minimal substrate contribution) scenarios; dashed lines refer to the s = 100 % (maximal substrate contribution) scenarios. See 4.2.1 and Table 3 for details of the conditions for each scenario.

**Figure 5.**
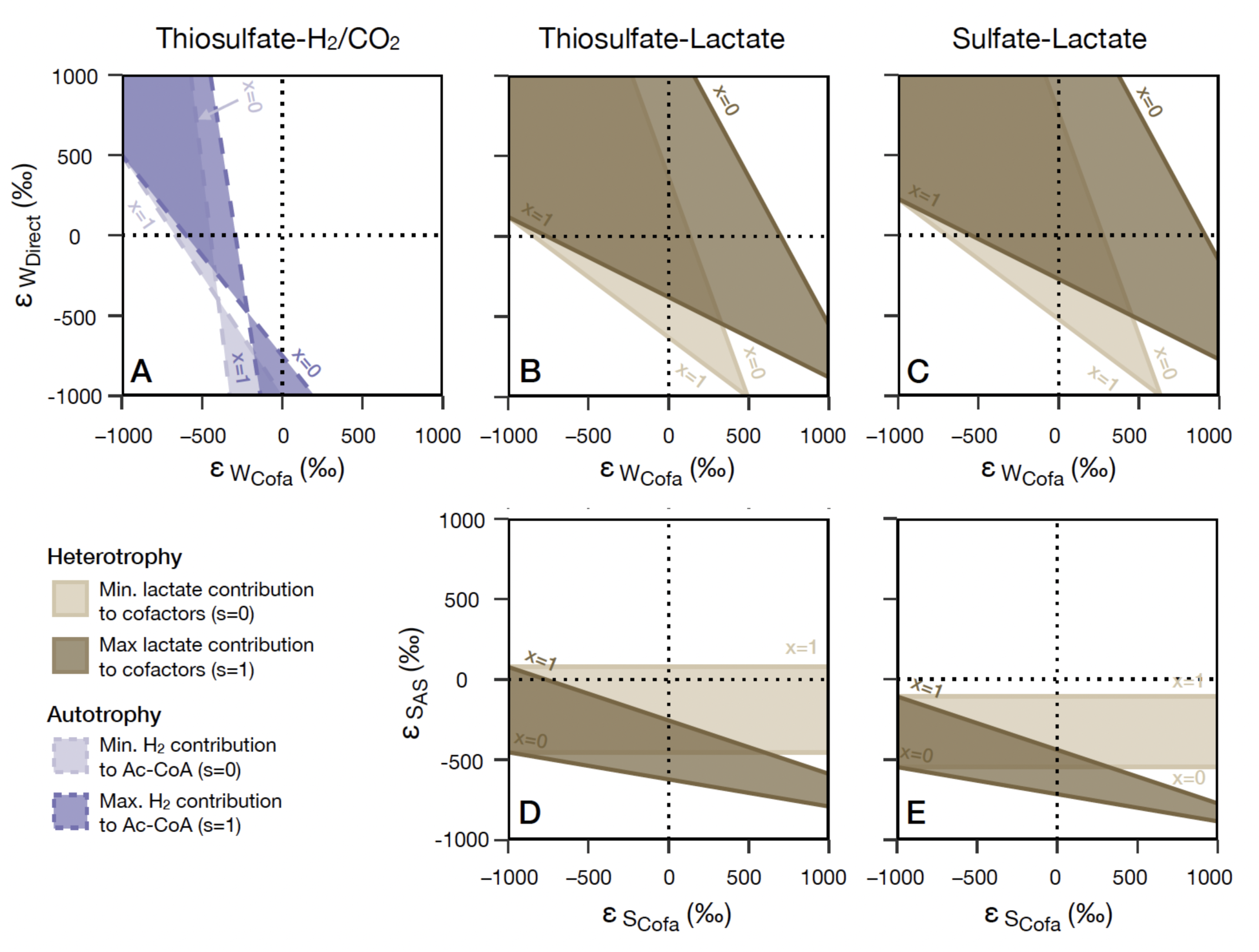
Isotope flux-balance model results for *Archaeoglobus fulgidus*. (A–C) Breakdown of the net isotope fractionation between lipid and water (ε_W,net_; Fig. 4D–F) into two different isotope fractionation factors: ε_W,direct_ for H atoms that are transferred through exchange during isomerization or addition via GGR, corresponding to *f*_W,direct_ in Fig. 3, and ε_W,cofa_ for H atoms that are transferred through cofactors (Fd_red_ and F_420_H_2_ for autotrophy, panel A; Fd_red_ for heterotrophy, panels B and C), corresponding to *f*_W,cofa_ in Fig. 3. (D–E) Breakdown of the net isotope fractionation between lipid and substrate (ε_S,net_; Fig. 4G–I) into two different isotope fractionation factors: ε_S,AS_ for H atoms that are inherited directly from substrate via Ac-CoA, corresponding to *f*_S,AS_ in Fig. 3, and ε_S,cofa_ for H atoms that are transferred through cofactors (only applicable for heterotrophy via F_420_H_2_, panels D and E), corresponding to *f*_S,cofa_ in Fig. 3. The lighter shaded areas refer to the s = 0 % (minimal substrate contribution) scenarios; darker shaded areas refer to the s = 100 % (maximal substrate contribution) scenarios. Solid lines and brown colors refer to heterotrophic conditions; dashed lines and purple colors refer to autotrophic conditions. See 4.2.1 and Table 3 for details of the conditions for each scenario.

For the net isotope fractionation between lipid and water (ε_W,net_), the two heterotrophic cases showed similar ranges and patterns where the ε_W,net_ values decrease with increasing *x*_ex_ (Fig. 4E– F). The ε_W,net_ range was relatively more positive during stationary growth phase (–218 to 776 ‰ for the T-L condition; –259 to 682 ‰ for the S-L condition) compared to exponential growth phase (–312 to 561 ‰ for the T-L condition; –319 to 546 ‰ for the S-L condition) (Table 3: Fig. 4E–F). For the autotrophic case, the growth phase-dependent trend was also observed (more positive ε_W,net_ during stationary phase; Table 3; Fig. 4D). Unlike in the heterotrophic cases, all ε_W,net_ values for the autotrophic case were negative and fell within a relatively narrow range for the entire range of *x*_ex_ (–448 to –312 ‰ for exponential phase; –372 to –228 ‰ for stationary phase; Table 3). For the autotrophic case, a theoretical end-member ε_W,net_ with no substrate contribution can be estimated by calculating the regression slope through the origin (i.e., no intercept, which represents the substrate contribution term, f_S_·^2^ɑ_L/W_·R_S_; Eq. (4)): –293 ‰ for the autotrophic case (black dotted line; Fig. 4D). This value falls within the range of ε_W,net_ for stationary, rather than exponential, growth phase (Fig. 4D). For the net isotope fractionation between lipid and substrate (ε_S,net_), the two heterotrophic cases showed similar ranges and patterns where ε_S,net_ values increase with increasing *x*_ex_ (Fig. 4H–I). The growth phase-dependent trend for ε_S,net_ contrasted that of ε_W,net_, where the ε_S,net_ range was relatively more negative during stationary growth phase (–688 to –282 ‰ for the T-L condition; –654 to –201 ‰ for the S-L condition) compared to exponential growth phase (–582 to –36 ‰ for the T-L condition; –592 to –59 ‰ for the S-L condition) (Table 3; Fig. 4H–I). A similar trend was observed for the autotrophic case (more negative ε_S,net_ during stationary phase; Table 3; Fig. 4G).

The net isotope fractionation factors (ε_W,net_ and ε_S,net_) can be further broken down into individual fluxes associated with H isotopes with different routes of incorporation. The net isotope fractionation factor between lipid and water (ε_W,net_) can be divided into: 1) ε_W,direct_ associated with H atoms that are transferred through the f_W,direct_ flux (Fig. 3) via H exchange between isoprenoid precursors and intracellular water or addition via GGR, and 2) ε_W,cofa_ associated with the f_W,cofa_ flux (f_W,cofa-LS_ for heterotrophy; f_W,cofa-LS_ and f_W,cofa-AS_ for autotrophy; Fig. 3) via cofactors (Fd_red_ and F_420_H_2_ for autotrophy; Fd_red_ for heterotrophy) (Fig. 5A–C). The net isotope fractionation factor between lipid and substrate (ε_S,net_) can be divided into: 1) ε_S,AS_ associated with H atoms that are transferred through the f_S,AS_ flux (Fig. 3) via Ac-CoA synthesis, and 2) ε_S,cofa_ associated with the f_S,cofa_ flux (f_S,cofa-NADPH_ and f_S,cofa-GGR_ for heterotrophy; Fig. 3) via cofactors (F_420_H_2_) (Fig. 5D–E). Interpretation of these results are discussed further in 5.2.1.

## 5. Discussion

### 5.1. Effects of carbon metabolism and growth phase on lipid H isotopes composition in Archaea

The isotope flux-balance model results show a large fraction of lipid-H directly reflects water H isotope compositions, rather than substrate (lactate or H_2_) isotope compositions in *A. fulgidus*. The *f*_W,net_ is the net fraction of total biphytane H that directly records water H isotopes and is explicitly not derived from organic substrate or H_2_. We observe larger *f*_W,net_ for autotrophic growth (*f*_W,net_ = 80–100 %), whereas *f*_W,net_ for heterotrophic growth was lower (*f*_W,net_ = 31–70 %; Table 3; Fig. 4A– C). Given that *ε*_W,net_ is likely negative (i.e., a normal kinetic isotope fractionation), we can constrain the *f*_W,net_ range that corresponds to ε_W,net_ < 0 (*see* Fig. 4D–E). This yields *f*_W,net_ ranges of: 38–52 % for T-L and 42–52 % for S-L during exponential growth phase, versus 50–56 % for T-L and 52–54 % S-L during stationary growth phase. The lowest *f*_W,net_ recorded were during heterotrophy in exponential phase, independent of the terminal electron acceptor. For stationary phase cultures of *A. fulgidus*, the *f*_W,net_ was ≥80 % for autotrophy versus ≥50 % for heterotrophy. The *f*_W,net_ range for lipids produced during stationary phase alone is likely larger, because lipids collected during stationary phase are a mixture of those produced during exponential phase and those newly produced during stationary phase. The higher *f*_W,net_ range observed for autotrophy is reasonable, given that lactate is an additional source of H that contributes directly to Ac-CoA synthesis during heterotrophy. A similar observation was reported for bacterial lipids where, for instance, the mole fraction of lipid H derived from water (≡ *X*_W_ in Zhang et al., 2009; herein equivalent to *f*_W,net_) was larger for *Escherichia coli* grown on glucose compared to the same organism grown on complex medium (Zhang et al., 2009). Though the lipid synthesis pathways are different for Bacteria versus Archaea, it appears that the fractional contribution of substrates that are more direct precursors for lipid biosynthesis generally increases when organic substrates are used or H_2_ is supplied in excess as an electron donor.

The effects of carbon metabolism and growth phase on lipid isotope composition have not been investigated in detail for archaeal lipids. Our results indicate that both the mode of metabolism and growth phase affect the lipid isotope composition (Table 3; Fig. 4). A recent study reported ε_L/W_ of BPs produced by *N. maritimus* in continuous cultures at a range of doubling time and observed little change in ε_W/L_ (0.18 ± 0.09 ‰ increase in ε_L/W_ per hour increase in doubling time; Leavitt, Kopf et al., 2023). The relative variation in doubling time we observe for *A. fulgidus* grown on different substrates here in batch cultures (Table 2) is similar (*ca*. 3-fold variation) to the variation observed for *N. maritimus* in Leavitt, Kopf et al. (2023). A key difference here is that we do not strictly control growth and metabolic rates, as was the case for chemostat cultivation of *N. maritimus* (Leavitt, Kopf et al., 2023), and for *Sulfolobus acidocaldarius* (Harris et al., 2022). Nonetheless, the range of ε_L/W_ we observe across different metabolic modes by *A. fulgidus* (–231 ± 6 ‰ for T-L, –241 ± 4 ‰ for S-L, and –271 ± 18 ‰ for T-H_2_/CO_2_; Table 2) is significant compared to the entire range of ε_L/W_ observed across a similar variation in growth rate by *N. maritimus* (–272 ± 20 to –279 ± 8 ‰ for BP-0; Leavitt, Kopf et al., 2023). *N. maritimus* is an obligate chemoautotroph, such that all BP-bound H is strictly biosynthetic, whereas we show here that patterns of ε_W/L_ can change between heterotrophic and autotrophic growth conditions (Fig. 4 and Fig. 6). Our results support this hypothesis and show variation in ε_L/W_ up to ∼40 ‰ within the same metabolically versatile organism, in response to different electron donor-acceptor pairs and modes of carbon metabolism.

**Figure 6.**
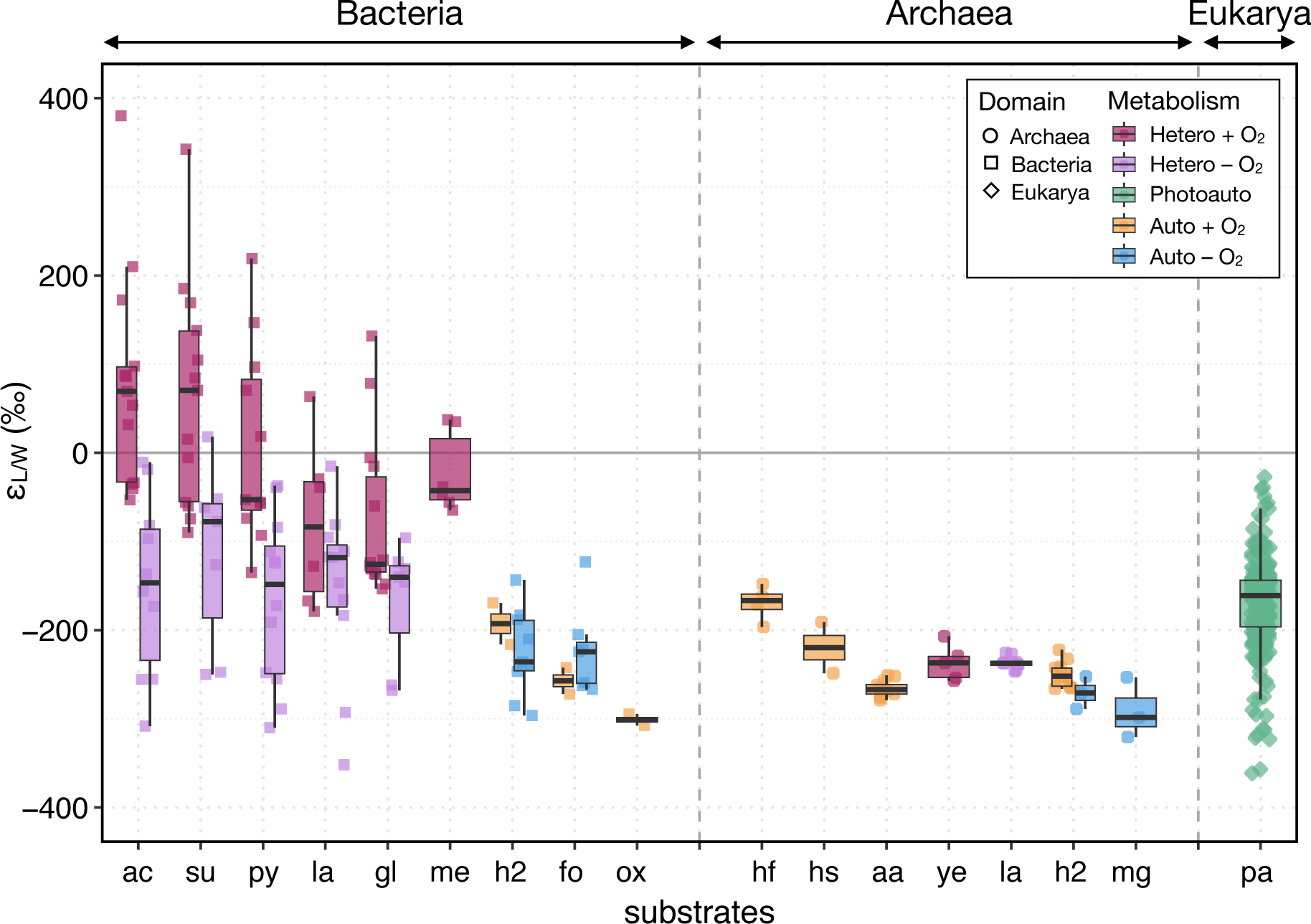
Summary of hydrogen isotope fractionation (ε_L/W_) between lipids and water observed in culture studies. ε_L/W_ values observed in all three domains of life: Archaea (circle), Bacteria (square), and Eukarya (diamond). Color codes represent different types of metabolism: aerobic heterotrophy (red), anaerobic heterotrophy (purple), oxygenic photoautotrophy (green), aerobic chemoautotrophy (orange), and anaerobic chemoautotrophy (blue). Electron donors used for biosynthesis are shown on the horizontal axis: acetate (ac); succinate (su); pyruvate (py); lactate (la); glucose (gl); methane (me); H_2_+CO_2_ (h2); formate (fo); oxalate (ox); H_2_+Fe^3+^ (hf); H_2_+S^0^ (hs); ammonium (aa); yeast extract (ye); H_2_, acetate or methanol for methanogenesis (mg); water for photoautotrophy (pa). Tested organisms for bacterial cultures include *C. oxalaticus*, *C. necator*, *R. palustris* (Zhang et al., 2009); *E. coli* (Zhang et al., 2009; Osburn et al., 2016; Wijker et al., 2019); *D. autotrophicum* (Campbell et al., 2009; Osburn et al., 2016); *M. capsulatus* (Sessions et al., 2002); *D. multivorans* (Dawson et al., 2015); *D. alaskensis* (Leavitt et al., 2016); *P. denitrificans*, *S. oneidensis*, *D. hydrogenophilus*, *D. alaskensis*, *D. propionicus* (Osburn et al., 2016); *B. subtilis*, *P. fluorescens*, *R. radiobacter*, and *E. meliloti* (Wijker et al., 2019). Tested organisms for eukaryotic cultures include *A. fundyense*, *I. galbana*, *Ascophyllum* sp., *F. vesiculosis*, *Z. marina*, *S. alterniflora* (Sessions et al., 1999); *C. japonica* (Chikaraishi et al., 2004); *B. braunii*, *E. unicocca*, *V. aureus* (Zhang and Sachs, 2007); *C. sativuc* (Chikaraishi et al., 2009); *E. huxleyi* (Sachs et al., 2016). Tested organisms for archaeal cultures include *Acidianus* sp. DS80 (this study); *N. maritimus* (Leavitt, Kopf et al., 2023); *Sulfolobus* sp. (Kaneko et al., 2011); *A. fulgidus* (this study); *Metallosphaera sedula* (this study); and *M. barkeri* (Wu et al., 2020).

Even though the range of ε_L/W_ in *A. fulgidus* is relatively larger compared to *N. maritimus*, the overall variation of ε_L/W_ across all archaeal lipid studies to-date is consistently small compared to the variation observed across bacterial lipids (Fig. 6).

### 5.2. Factors that affect lipid-water H isotope fractionation across the three domains of life

Here we present a synthesis of the broader ε_L/W_ patterns observed in lipids from all three domains of life. Archaeal lipids are characterized by a relatively narrow range of large and negative values of ε_L/W_ across a wide range of organisms, metabolisms, and growth conditions (Fig. 6; Kaneko et al., 2011; Wu et al., 2020; Leavitt, Kopf et al., 2023; this study). This is in contrast with the domain Bacteria, where fatty acid ε_L/W_ ranges a large span of >600 ‰ with both highly positive and negative values (Fig. 6; Campbell et al., 2009; Zhang et al., 2009; Dawson et al., 2015; Osburn et al., 2016; Wijker et al., 2019). Archaeal lipid ε_L/W_ largely overlaps with that of eukaryotic lipid ε_L/W_, although eukaryotic lipid ε_L/W_ spans a significant range depending on the types of organism and classes of compound (Fig. 6; Sessions et al., 1999; Chikaraishi et al., 2004, 2009; Zhang and Sachs, 2007; Sachs et al., 2016). Notably, there are subgroups of both domains Bacteria and Eukarya that share similarities with Archaea in their ε_L/W_. The shared similarities are interesting, given the difference in lipid biosynthesis mechanism among the three domains. Below, we take a closer look at the specific subsets of ε_L/W_ values from the domains Euakrya and Bacteria that resemble those from Archaea and discuss factors that may contribute to the large, negative values of ε_L/W_ observed across the three domains.

#### 5.2.1. Large lipid-water fractionation upon isoprenoid saturation in Archaea and Eukarya

The consistently large, negative values of ε_W/L_ for archaeal lipids are notable considering the different types of organism and metabolic modes studied so far (Fig. 6). In Fig. 5, we present the model solution spaces for individual H fluxes, within either net flux that reflects water isotope composition (i.e., not affected by substrate contributions; *f*_W,net_) or net flux that reflects isotopic contributions from organic substrate or H_2_ (*f*_S,net_), to further investigate specific parts of archaeal lipid biosynthesis pathway that may contribute to the observed large and negative values of ε_L/W_. Prior work yielded estimates of α_W_ ≅ 0.9 for isotope fractionation associated with direct water incorporation across many bacteria (Zhang et al., 2009; Wijker et al., 2019), and ε_L/W_ values modeled for *N. maritimus* with this empirical α_W_ were consistent with experimental observations (Leavitt, Kopf et al., 2023). Assuming an equivalent value of ε_W,direct_ (i.e., isotope fractionation associated with the *f*_W,direct_ flux is –100 ‰), we constrain plausible ranges of ε_W,cofa_ in *A. fulgidus* (Fig. 5A–C). The ranges of ε_W,cofa_ at ε_W,direct_ = –100 ‰ are particularly well constrained for the autotrophic case (–599 to –436 ‰ at *s* = 0 %; –519 to –276 ‰ at *s* = 100 %; Fig. 5A).

It is plausible that both ε_W,cofa-AS_ and ε_W,cofa-LS_ have negative values that are comparable to the combined value of ε_W,cofa_, as the less likely alternative would be a negative ε_W,cofa_ that consists of very different ε_W,cofa-AS_ and ε_W,cofa-LS_. Our current model cannot distinguish between isotope fractionation factors associated with different fluxes within *f*_W,cofa_ (i.e., *f*_W,cofa-AS_ through Ac-CoA *vs*. *f*_W,cofa-LS_ through NADPH), both of which can inherit H from cofactors F_420_H_2_ and/or Fd_red_ (Fig. 3B). However it is worth noting that, for both heterotrophic and autotrophic cases, a fraction of the *f*_W,cofa-LS_ flux represents hydride transfer during the final saturation (double bond reduction) via GGR (Fig. 3; Step 10, Appendix B). Results of the isotope flux-balance model for *N. maritimus* also indicated that the GGR reduction step is strongly fractionating (ε_GGR_ = –690 to –820 ‰ if ε_NADPH_ = –100 to –300 ‰, depending on the degree of water exchange, or *x*_ex_ in this study), regardless of how other model parameters are set (Leavitt, Kopf et al., 2023). Given how *N. maritimus* and *A. fulgidus* share little in common otherwise (metabolically, genetically, and biochemically), large fractionation associated with the GGR reduction step provides a parsimonious explanation for the similar ε_L/W_ values observed in the two organisms, as it is the final step of archaeal lipid biosynthesis shared across all tetraether-producing Archaea. This would be consistent with the similarly large, negative values of ε_L/W_ observed in other archaeal lipids (Fig. 6); testing more Archaea with the combined approach of isotope labeling experiments and biochemically informed isotope flux-balance model will help further confirm this idea.

A closer examination of the eukaryotic lipids that share similar ε_L/W_ values to archaeal lipids lends further support to the notion that GGR reduction contributes to large H isotope fractionation between lipids and water. Fig. 7A shows ε_L/W_ for different subgroups of eukaryotic lipids, categorized by three different biosynthetic pathways. In general, the two major families of lipids are acetogenic and isoprenoid lipids which consist of acetate and isoprene monomeric units, respectively (Pearson, 2014). Isoprenoid lipids are further divided into two groups, depending on whether the isoprene (C_5_) building blocks are synthesized via the mevalonic acid (MVA) pathway (Lynen et al., 1958; Bloch, 1959) or the methylerythritol phosphate (MEP) pathway (Rohmer et al., 1993). All Archaea discussed in this study synthesize isoprenoid lipids via the MVA pathway. The MEP pathway is generally defined as bacterial and the MVA pathway as eukaryotic and archaeal; both pathways often simultaneously occur in plants, where the MEP pathway is expressed in chloroplasts (relics of the cyonobacterial plastid ancestor) and the MVA pathway is expressed in the cytosol (Lichtenthaler et al., 1997; Lichtenthaler, 1999; Pearson, 2014). An exception to this is the case of green algae, where both the cytosolic sterols and plastidic phytol isoprenoids are synthesized via the MEP pathway (Lichtenthaler et al., 1997; Disch et al., 1998; Pearson, 2014).

**Figure 7.**
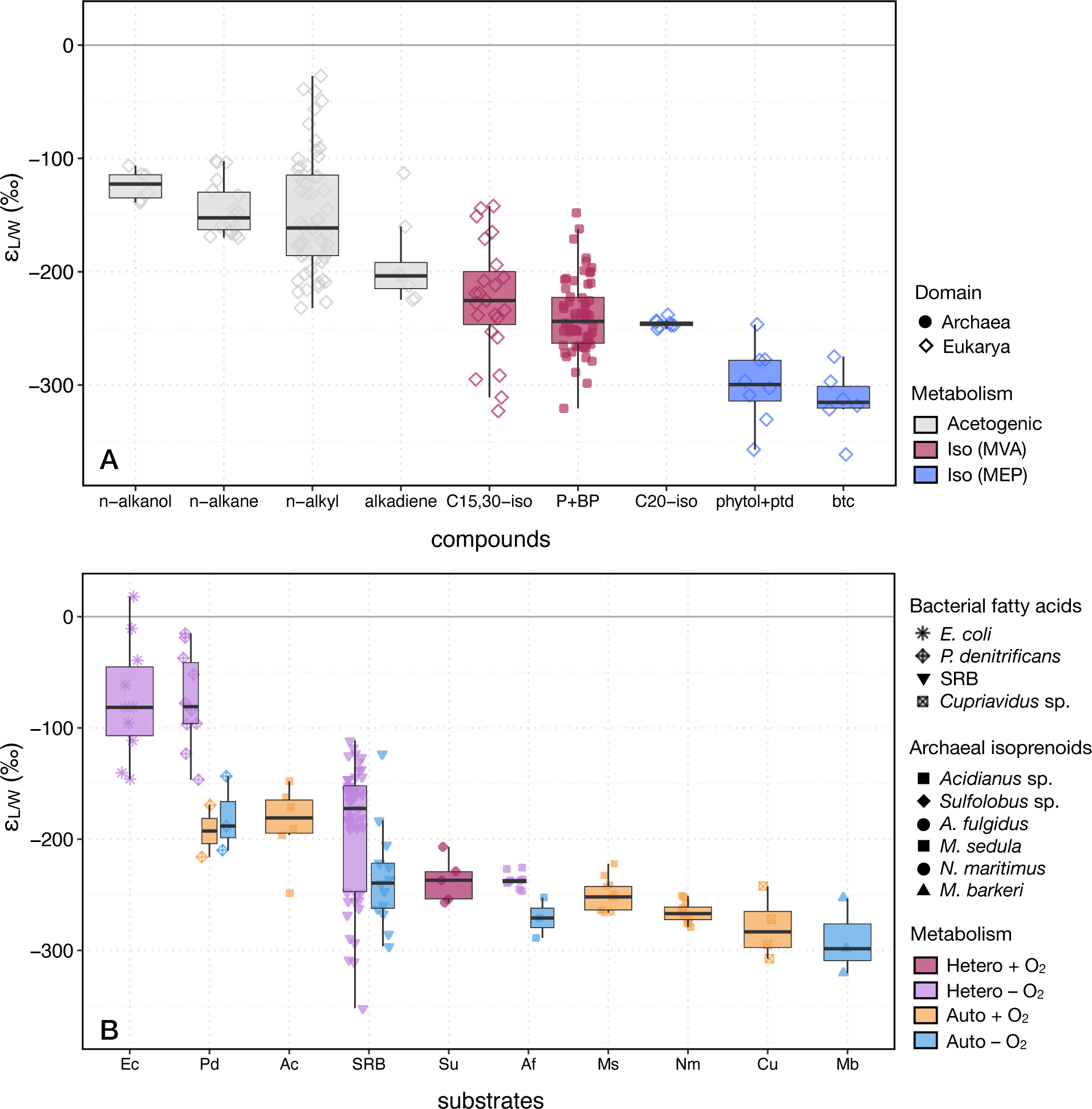
Comparison of ε_L/W_ values observed in eukaryotic *vs*. archaeal lipids and in bacterial *vs*. archaeal lipids. **(A)** ε_L/W_ values observed in acetogenic and isoprenoid lipids from the domains Archaea and Eukarya. Different classes of compounds are shown on the horizontal axis. Compounds shown for acetogenic pathway: n-alkanol, *n*-alkane, *n*-alkyl (*n*-alkanoic acids and fatty acids), and alkadiene. For the MVA pathway: C_15_ and C_30_ isoprenoids (sterol, squalene, triterpenoids, and sesquiterpenes) and P+BP (phytanes and biphytanes). For the MEP pathway: C_20_ isoprenoids (diterpenes, sandaracopimarinol, ferruginol, diterpenol, and sugiol), phytol+ptd (phytol and phytadiene), and btc (botryococcene). **(B)** ε_L/W_ values observed in fatty acids from anaerobic and/or autotrophic Bacteria and isoprenoid lipids from Archaea. Different organisms are shown on the horizontal axis. For Bacteria: *E. coli* (Ec), *Paracoccus denitrificans* (Pd), SRB (*Desulfobacterium autotrophicum*, *Desulfococcus multivorans*, *Desulfobacter hydrogenophilus*, *Desulfovibrio alaskensis*, *Desulfolobus propionicus*), and *Cupriavidus* sp. (Cu). For Archaea: *Acidianus* sp. DS80 (Ac), *Sulfolobus* sp. (Su), *Archaeoglobus fulgidus* (Af), *Metallosphaera sedula* (Ms), *Nitrosopumilus maritimus* (Nm), and *Methanosarcina barkeri* (Mb). A full list of references can be found in Appendix C.

The three different biosynthetic pathways of eukaryotic lipids (acetogenic, MVA, and MEP) are associated with distinct ranges of ε_L/W_. In general, isoprenoid lipids have more negative ε_L/W_ compared to acetogenic lipids (Estep and Hoering, 1980; Sessions et al., 1999). Among isoprenoid lipids, plastidic isoprenoids such as phytols are more depleted in ^2^H than the cytosolic counterparts (Sessions et al., 1999; Chikaraishi et al., 2004, 2009). This general trend is captured in Fig. 7A, where acetogenic lipids (e.g., *n*-alkanol and *n*-alkyl lipids including fatty acids) have more positive ε_L/W_ compared to isoprenoid lipids. The C_15_ and C_30_ isoprenoids synthesized via the MVA pathway (e.g., sterol, triterpenoids, sesquiterpenes, squalene) are relatively more enriched in ^2^H compared to the C_20_ isoprenoids (e.g., phytol, phytadiene, diterpenes) and botryococcenes (triterpenes) synthesized via the MEP pathway (Fig. 7A).

While the relative difference in ε_L/W_ between MVA and MEP pathways holds in general, the larger and more negative values of ε_L/W_ appear to correlate more with highly fractionating enzymatic steps, rather than specific pathways. For example, Chikaraishi et al. (2004) observed in *Cryptomeria japonica* (conifer) that ε_L/W_ values for cytosolic lipids produced via the MVA pathway, sesquiterpenes (–228 to –238 ‰) and squalene (–225 to –251 ‰), are similar to those for plastidic diterpenoids (–238 to –251 ‰) produced via the MEP pathway (Fig. 7A). Phytol and botryococcenes, on the other hand, have particularly negative ε_L/W_ values among eukaryotic lipids produced via the MEP pathway. For example, Zhang and Sachs (2007) found that botryococcenes with longer chain lengths had progressively lower δ^2^H compared to the C_30_ botryococcene (precursor to all other botryococcenes) in *Botryococcus braunii* race B. They hypothesized that significantly ^2^H-depleted hydrogen from methionine is incorporated into botryococcene during the step-wise methylation (Zhang and Sachs, 2007). Similarly, phytol in *C. japonica* had up to 65 ‰ more negative ε_L/W_ compared to diterpenoids, another plastidic lipid produced via the MEP pathway within the same organism (Chikaraishi et al., 2004). Later, Chikaraishi et al. (2009) observed a gradual depletion in δ^2^H from phytol precursors (starting from geranylgeraniol) to phytol in cucumber, indicating that significantly ^2^H-depleted hydrogen is incorporated into phytol and its precursors during the step-wise hydrogenation catalyzed by GGR (ε_GGR_ ≅ –650 ‰; Chikaraishi et al., 2009).

The ε_L/W_ values of phytol (and its dehydration product phytadiene) investigated in a wide range of eukaryotes so far are all significantly large and negative (e.g., –357 to –278 ‰ for phytol in dinoflagellate, coccolithophore, marsh grass, cucumber, and conifer; Fig. 7A). Similarly, all values of ε_L/W_ investigated in a range of different archaea are significantly large and negative (e.g., –341 to –148 ‰ for phytanes and biphytanes in *N. maritimus*, *Sulfolobus* sp., *M. sedula*, *Acidianus* sp., *M. barkeri*, and *A. fulgidus*; Fig. 7A). Taken together, empirical and model-based results suggest that the GGR reduction is a highly fractionating step that contribute to the large fractionation observed in both eukaryotic and archaeal lipid synthesis (Chikaraishi et al., 2009; Leavitt, Kopf et al., 2023; this study, Fig. 5 and Fig. 6).

#### 5.2.2. Energy limitation and lipid-water fractionation in Archaeal versus Bacterial lipids

Besides the shared biochemical mechanism underlying negative ε_L/W_ in eukaryotic and archaeal lipids, another interesting pattern emerges from comparison of ε_L/W_ across the three domains of life: a subset of bacterial lipids resembles archaeal lipids in its consistently negative ε_L/W_ values (Fig. 6). For bacterial lipids, anaerobiosis and/or autotrophy appear to be associated with consistently large, negative values of ε_L/W_. Examples include aerobic autotrophy (*Cupriavidus oxalaticus*, Zhang et al., 2009; *Paracoccus denitrificans*, Osburn et al., 2016), anaerobic autotrophy (*Desulfobacterium autotrophicum*, Campbell et al., 2009; *Desulfococcus multivorans*, Dawson et al., 2015; *P. denitrificans*, *Desulfobacterium autotrophicum*, *Desulfovibrio alaskensis*, Osburn et al., 2016), and anaerobic heterotrophy (*D. multivorans*, Dawson et al., 2015; *P. denitrificans*, *Escherichia coli*, *D. autotrophicum*, *Desulfobacter hydrogenophilus*, *D. alaskensis*, *Desulfolobus propionicus*, Osburn et al., 2016) (Fig. 6; Fig. 7B). Of these modes of metabolism, aerobic or anaerobic autotrophy has a more constrained range of ε_W/L_ and more overlaps with ε_L/W_ observed in archaeal lipids, compared to anaerobic heterotrophy (Fig. 6; Fig. 7B). We propose that NADPH flux imbalances associated with energy limitation that broadly characterize autotrophic and/or anaerobic modes of metabolism contribute to the apparent similarity in ε_L/W_ observed between the domains of Bacteria and Archaea.

The concept of NADPH flux imbalance set forth by Wijker et al. (2019) provides a quantitative framework to relate metabolism-dependent patterns of ε_L/W_ observed in aerobic heterotrophs to the fluxes of H through specific dehydrogenase and transhydrogenase enzymes. For the highly conserved fatty acid biosynthesis pathway, NADPH serves as the reducing agent for the reduction of acyl intermediates (Wakil et al., 1983). As a result, NADPH accounts for ∼50% of fatty acid H (Sessions et al., 1999). Wijker et al. (2019) showed that ε_L/W_ values of fatty acids produced by five wild-type species of aerobic heterotrophic bacteria are positively correlated with relative NADPH imbalance flux. Among the tested aerobic heterotrophs, two species (*E. coli* and *B. subtilis*) had moderately negative values of NADPH flux imbalance (–32 and –11 %, respectively) (Wijker et al., 2019). Here, negative values reflect underproduction of NADPH where catabolic production of NADPH does not meet the anabolic requirement (Wijker et al., 2019). Furthermore, the results from *E. coli* knockout mutants confirmed the direction of isotope fractionation for specific transhydrogenases (e.g., ^2^H-depletion by NADPH-producing PntAB and ^2^H-enrichment by NADPH-consuming UDhA) (Wijker et al., 2019).

For microbes adapted to chronic energy limitation, even more negative values of NADPH flux imbalance would be expected where just enough NADPH is produced such that most to all of it is used for biosynthesis (i.e., little to no flux from NADPH back to NADH). This prediction is consistent with the general concept of energy spilling (also known as growth uncoupling or overflow metabolism), which suggests that microbes under energy limitation tend to have more efficient biomass production (Russell, 2007). For example, *E. coli* grown aerobically resulted in more than 9-fold increase in ATP production per mole of glucose compared to the anaerobic counterpart; however, the biomass yield was only 2-fold greater during aerobic respiration (Stouthamer and Bettenhaussen, 1977; Gottschalk, 1986). Similar phenomenon of energy spilling has also been observed in methanogenic archaea. For example, *Methanothermobacter thermautotrophicus* and *Methanococcus maripaludis* produced less biomass per mole of methane when H_2_ (electron donor) was present in excess compared to H_2_-limiting condition (Morgan et al., 1997; De Poorter et al., 2007; Costa et al., 2013).

In line with the observed and expected outcomes of energy limitation (negative NADPH flux imbalance and tighter coupling between catabolism and anabolism), Leavitt, Kopf et al. (2023) found that ε_L/W_ values for *N. maritimus* fall within the 95% confidence interval for the extension of Wijker et al. (2019) regression when an extremely negative value (–100 %) of NADPH flux imbalance was assigned. The apparent fit of *N. maritimus* ε_L/W_ on the negative end of the framework is reasonable, given that *N. maritimus* is an obligate autotroph adapted to oligotrophic nitrification with energy-efficient CO_2_ fixation pathway (Martens-Habbena et al., 2009; Könneke et al., 2014). The fact that all archaeal lipids investigated for ε_L/W_ so far fall within a relatively narrow range of negative values (Fig. 6; Fig. 7B) may, at least partially, reflect the general concept that Archaea are optimized for chronic energy limitation (Valentine, 2007). The results from this study show that, while metabolic modes and growth rates have significant impacts on ε_L/W_ within a single organism (*A. fulgidus*), the observed variation in ε_L/W_ still falls within the overall range observed across archaeal lipids (e.g., heterotrophic and autotrophic *A. fulgidus* ‘Af’ data *vs*. other archaeal lipids data; Fig. 7B).

The similarity between archaeal lipids and a subset of anaerobic and/or autotrophic bacterial lipids is striking (Fig. 6; Fig. 7B) given the fundamental differences in fatty acid and isoprenoid lipid biosynthesis pathways. This suggests that factors controlling microbial ε_L/W_ may not be domain-specific; instead, common challenges associated with anaerobic respiration and/or autotrophy (i.e., lower free energy yields and/or additional energetic costs for CO_2_ fixation) may become primary controlling factors of ε_L/W_ for both Bacteria and Archaea. Notably, the range of ε_L/W_ in fatty acids produced by sulfate-reducing bacteria (SRB) is much narrower compared to that observed in aerobic heterotrophs regardless of metabolic pathway or substrate (Fig. 7B).

One of the transhydrogenases that plays an important role in SRB is NfnAB, and it has been shown that this enzyme is important for H isotope fractionation in *D. alaskensis* (Leavitt et al., 2016). The details of the magnitude of ε_L/W_ set by NfnAB under severe NADPH underproduction may be complicated by other processes such as intracellular isotope distillation (i.e., more quantitative transfer of NADPH hydride to lipids) as suggested by Leavitt et al. (2016). Regardless, given that decoupling of NADPH production for anabolism from NADH for catabolism becomes more important during energy limitation, transhydrogenases that are widely distributed among anaerobic bacteria (namely NfnAB; Pereira et al., 2011; Leavitt et al., 2016; Buckel and Thauer, 2018) may be key contributing factors for the negative ε_W/L_ observed across these organisms (Fig. 6; Fig. 7B). It is worth noting that *nfnAB* was not found in archaeal species investigated by Pereira et al. (2011), including *A. fulgidus*.

A complex interplay among environmental, physiological, and enzymatic factors underlies net isotope fractionation patterns observed in a biosynthetic product such as ε_L/W_. For archaeal lipids and a subset of bacterial lipids, however, two factors appear to play key roles in shaping the consistently negative ε_L/W_ patterns observed in the lipids from both domains: highly fractionating enzymatic steps (e.g., GGR in archaeal isoprenoid lipid biosynthesis; transhydrogenases in bacterial fatty acid biosynthesis) and energy limitation (i.e., NADPH flux imbalance). The negative ε_L/W_ commonly observed in archaea and a subset of bacteria is likely a convergent feature resulting from a combination of both factors. A better mechanistic understanding of factors conducive to similar patterns of ε_L/W_ in bacterial and archaeal lipids could lead to biogeochemical and/or ecological insights that are unique to microbial life. This would, in turn, significantly expand the application of lipid biomarkers as paleoenvironmental or paleoecological proxy currently possible with eukaryotic or bacterial lipids alone.

## 6. Conclusion

In this study, we conducted water isotope label experiments with a metabolically flexible and well-studied model archaeon *A. fulgidus* and quantified the hydrogen isotope fractionation between lipids and water in response to different carbon substrates and electron donor-acceptor pairs. We observed changes in regression parameters among experimental conditions, where higher slopes (*f*_W_·^2^α_L/W_) were observed in autotrophic compared to heterotrophic conditions and during stationary compared to exponential growth phase. The bio-isotopic model further constrains individual parameters such as fluxes that reflect water isotope composition (i.e., not affected by substrate isotope composition) (*f*_W_) and isotope fractionation between lipids and water (ε_L/W_) that together make up the slope. These results are consistent with observations from regression parameters, where *f*_W_ is larger under autotrophic condition and during stationary growth phase compared to the heterotrophic conditions and exponential growth phase. The model results also allow us to constrain plausible ranges of isotope fractionation associated with a specific route of hydrogen flux (e.g., flux for direct water incorporation *vs*. flux from water through cofactors). The model results point toward the isotope fractionation associated with the *f*_W,cofa_ flux, which includes the hydride transfer during the final saturation via GGR, as a highly fractionating step. This is consistent with recent bio-isotopic model results for another archaeon *N. maritimus*. Finally, we synthesized a three-domain comparison of ε_L/W_ patterns which suggests that the highly fractionating double bond reduction via GGR may also explain the similarly large and negative values of ε_L/W_ observed between eukaryotic and archaeal lipids. Furthermore, the shared negative ε_L/W_ values between fatty acids in anaerobic and/or autotrophic bacteria and isoprenoids in archaea may be associated with the general state of energy limitation experienced by the different groups of organisms. Additional experimental work with archaea that use carbon metabolism pathways that have not been examined with the bio-isotopic model (i.e., 3HP/4HB and reductive Wood-Ljungdahl pathways) as well as more systematic comparisons between bacterial and archaeal lipids will help determine the scope of applicability for microbial lipid ε_L/W_ in paleoenvironmental and paleoecological studies.

## 7. Data availability

Access to all data products and code for the isotope flux-balance model for this manuscript are available at: https://github.com/jrhim/Rhim_et_al_2024_GCA.

## 8. CRediT authorship contribution statement

**Jeemin H. Rhim:** Conceptualization, Methodology, Investigation, Data Curation, Writing – Original Draft, Writing – Review & Editing, Visualization. **Sebastian Kopf:** Conceptualization, Methodology, Resources, Writing – Review & Editing, Supervision, Funding acquisition. **Jamie McFarlin:** Investigation, Writing – Review & Editing. **Ashley E. Maloney:** Investigation, Writing – Review & Editing. **Harpreet Batther:** Investigation, Writing – Review & Editing. **Carolynn M. Harris:** Investigation, Writing – Review & Editing. **Alice Zhou:** Investigation, Writing – Review & Editing. **Xiahong Feng:** Investigation, Writing – Review & Editing. **Yuki Weber:** Investigation, Writing – Review & Editing. **Shelley Hoeft-McCann:** Investigation, Writing – Review & Editing. **Ann Pearson:** Resources, Writing – Review & Editing, Supervision, Funding acquisition. **William D. Leavitt:** Conceptualization, Resources, Writing – Review & Editing, Supervision, Funding acquisition.

## 9. Declaration of competing interest

The authors declare that they have no known competing financial interests or personal relationships that could have appeared to influence the work reported in this paper.

## 10. Acknowledgements

This research was supported by funding from: collaborative research grant NSF EAR #1928303 (WDL, SHK); the American Chemical Society PRF #57209-DNI2 (WDL, YW); Simons Foundation Award #623881 (WDL); Swiss National Science Foundation P2BSP2_168716 (YW); NSF OCE-1843285 and 1702262 (AP). JHR was supported by the Dartmouth College Society of Fellows. We thank Ashley Maloney (CU Boulder), Felix Elling (then at Harvard), and Beverly Chiu (then at Dartmouth) for expert laboratory assistance.

## 11. Appendix A. Supplementary material

These include the Overview of hydrogen budget, Culturing conditions for *Acidianus* sp. And *Metallosphaera sedula*, Full list of references for Figure 7, and Supplemental Figures 1–2.

## Notes

### Competing Interest Statement

The authors have declared no competing interest.

### Summary of Updates

This version of the manuscript is the final version submitted to a journal for peer review. Updates include some formatting changes, minor figure and figure caption updates, and minor re-wording in the main text. One additional author (A. Maloney) has been added to the author list.

