## Supplementary Material for "Metabolic imprints in the hydrogen isotopes of *Archaeoglobus fulgidus* tetraether lipids"

### Overview of hydrogen budget

In our model for *A. fulgidus*, we consider two routes of H incorporation into lipids for both heterotrophic and autotrophic model cases: 1) via organic substrate or $H_2$ and 2) without any isotopic contribution from the substrates. Substrate refers to lactate and $H_2$, respectively, for the heterotrophic (Fig. 3A) and autotrophic cases (Fig. 3B). The stoichiometric budget for H sources in *A. fulgidus* can be simplified with the following fractions that make up either of the two passages for water H incorporation and substrate H incorporation. For the 80 H atoms in BP produced via heterotrophy ($BP_{Het}$): H incorporated directly via lipid biosynthesis ($f_{W,LS}$) and GGR reduction ($f_{W,GGR}$) are combined to yield total direct water H ($f_{W,direct}$); H incorporated indirectly via reducing cofactors NADPH ($f_{W,Cofa\text{-}NADPH}$) and GGR ($f_{W,Cofa\text{-}GGR}$) are combined to yield total indirect water H during lipid synthesis ($f_{W,Cofa\text{-}LS}$); H incorporated indirectly via lactate (substrate) and reducing cofactors NADPH ($f_{S,Cofa\text{-}NADPH}$) or GGR ($f_{S,Cofa\text{-}GGR}$) are combined to yield total indirect water H during lipid synthesis ($f_{S,Cofa\text{-}LS}$); and H incorporated directly via lactate during Ac-CoA synthesis ($f_{S,AS}$).

$$BP_{Het} = f_{W,direct} + f_{W,Cofa\text{-}LS} + f_{S,Cofa\text{-}LS} + f_{S,AS}$$
$$BP_{Het} = (f_{W,LS} + f_{W,GGR}) + (f_{W,Cofa\text{-}NADPH} + f_{W,Cofa\text{-}GGR}) + (f_{S,Cofa\text{-}NADPH} + f_{S,Cofa\text{-}GGR}) + f_{S,AS}$$
Eq. (A5)

For the 80 H atoms in BP produced via autotrophy ($BP_{Auto}$): H incorporated directly via lipid biosynthesis ($f_{W,LS}$) and GGR reduction ($f_{W,GGR}$) are combined to yield total direct water H ($f_{W,direct}$); H incorporated indirectly via reducing cofactors NADPH ($f_{W,Cofa\text{-}NADPH}$) and GGR ($f_{W,Cofa\text{-}GGR}$) are combined to yield total indirect water H during lipid synthesis ($f_{W,Cofa\text{-}LS}$); H incorporately indirectly via reducing cofactors $Fd_{red}$ ($f_{W,Cofa\text{-}Fdred}$) and $F_{420}H_2$ ($f_{W,Cofa\text{-}F420H2}$) are combined to yield total indirect water H during Ac-CoA synthesis ($f_{W,Cofa\text{-}AS}$); and H incorporated directly via $H_2$ (substrate) during Ac-CoA synthesis ($f_{S,AS}$).

$$BP_{Auto} = f_{W,direct} + f_{W,Cofa\text{-}LS} + f_{W,Cofa\text{-}AS} + f_{S,AS}$$
$$BP_{Auto} = (f_{W,LS} + f_{W,GGR}) + (f_{W,Cofa\text{-}NADPH} + f_{W,Cofa\text{-}GGR}) + (f_{W,Cofa\text{-}Fdred} + f_{W,Cofa\text{-}F420H2}) + f_{S,AS}$$
Eq. (A6)

These fractions represent the relative fluxes of H (i.e., individual arrows either directly or indirectly connecting the two ultimate H sources, water and substrate, to lipids) shown in Fig. 3. We explore four scenarios for each case to explore the degree of water exchange, $x_{ex}$, ranging from 0% of minimal exchange to 100% of maximal exchange and the degree of substrate contribution, ranging from 0% of minimal contribution to 100% of maximal contribution. For the heterotrophic case, the assigned fractions are as follows (Eq. A7):

$$f_{W,LS} = \left[\frac{2}{3} \cdot (1 - x_{ex}) + 24 \cdot x_{ex}\right] \div 80$$

$$f_{W,GGR} = 8 \div 80$$
$$f_{W,direct} = \left[\frac{2}{3} \cdot (1 - x_{ex}) + 24 \cdot x_{ex} + 8\right] \div 80$$
$$f_{W,cofa-NADPH} = \left[16 \cdot (1 - \frac{1}{3} \cdot s)\right] \div 80$$
$$f_{W,cofa-GGR} = \left[8 \cdot (1 - \frac{1}{3} \cdot s)\right] \div 80$$
$$f_{W,cofa-LS} = \left[(16 + 8) \cdot (1 - \frac{1}{3} \cdot s)\right] \div 80$$
$$f_{S,cofa-NADPH} = \left[16 \cdot \frac{1}{3} \cdot s\right] \div 80$$
$$f_{S,cofa-GGR} = \left[8 \cdot \frac{1}{3} \cdot s\right] \div 80$$
$$f_{S,cofa-LS} = \left[(16 + 8) \cdot (\frac{1}{3} \cdot s)\right] \div 80$$
$$f_{S,AS} = \left[24 + \left(23 + \frac{1}{3}\right) \cdot (1 - x_{ex})\right] \div 80$$

Eq. (A7)

where $x_{ex}$ is the degree of water exchange during lipid synthesis, which depends on the extent of re-equilibration during isomerization steps (steps 2 and 6; Appendix B). The $s$ term is the degree of H contribution to lipids from substrate (lactate). Fractions that contain the $x_{ex}$ term ($f_{W,direct}$ and $f_{S,AS}$; Eq. A7) vary between the two end-member scenarios of $x_{ex}$ = 0 % (minimal exchange) and $x_{ex}$ = 100 % (maximal exchange); fractions that contain the $s$ term ($f_{W,cofa-LS}$ and $f_{S,cofa-LS}$; Eq. A7) vary between the two end-member scenarios of $s$ = 0 % (minimal contribution) and $s$ = 100 % (maximal contribution). At maximal substrate contribution ($s$ = 100 %), ⅓ of $f_{W,cofa-LS}$ and $f_{S,cofa-LS}$ fractions are incorporated via substrates. During the complete oxidation of lactate to $CO_2$, the average carbon oxidation state increases from 0 to +4 (i.e., 12 electrons transferred total for all three carbons per molecule of lactate). The electron acceptors for the six oxidation steps are: menaquinone, for lactate to pyruvate oxidation (Step 1; Fig. 2A); ferredoxin, for pyruvate to Ac-CoA oxidation and the final two oxidation steps to $CO_2$ (Steps 3, 8 and 9; Fig. 2A); and $F_{420}$, for methenyl-$H_4$MPT and methylene-$H_4$MPT oxidation (Steps 4 and 5; Fig. 2A) (Möller-Zinkhan et al., 1989; Schmitz et al., 1991; Klein et al., 1993; Kunow et al., 1993; Schwörer et al., 1993). None of these redox reactions during lactate oxidation involve NAD or NADP as electron acceptors; in *A. fulgidus*, reduced pyridine nucleotides (NADPH) required for biosynthesis is produced via the reversible reduction of NADP with $F_{420}H_2$, similar to the case in methanogens (Kunow et al., 1993). Therefore, in the maximal substrate contribution scenario ($s$ = 100 %), up to ⅓ of H fluxes through the cofactor NADPH and GGR can be incorporated into NADP via the two (out of six, hence ⅓) hydrides in $F_{420}H_2$ via lactate ($f_{S,cofa-LS}$; Fig. 3A).

For the autotrophic case, the assigned fractions are as follows (Eq. A8):

$$f_{W,LS} = \left[\frac{2}{3}\cdot(1-x_{ex}) + 24\cdot x_{ex}\right] \div 80$$
$$f_{W,GGR} = 8 \div 80$$
$$f_{W,direct} = \left[\frac{2}{3}\cdot(1-x_{ex}) + 24\cdot x_{ex} + 8\right] \div 80$$
$$f_{W,Cofa-NADPH} = 16 \div 80$$
$$f_{W,Cofa-GGR} = 8 \div 80$$
$$f_{W,Cofa-LS} = (16+8) \div 80$$
$$f_{W,Cofa-Fdred} = \frac{1}{3}\cdot\left[24 + \left(23+\frac{1}{3}\right)\cdot(1-x_{ex})\right] \div 80$$
$$f_{W,Cofa-F420H2} = \left[\frac{1}{3}\cdot\left[24 + \left(23+\frac{1}{3}\right)\cdot(1-x_{ex})\right]\right]\cdot(2-s) \div 80$$
$$f_{W,Cofa-AS} = \left[\frac{1}{3}\cdot\left[24 + \left(23+\frac{1}{3}\right)\cdot(1-x_{ex})\right] + \left[\frac{1}{3}\cdot\left[24 + \left(23+\frac{1}{3}\right)\cdot(1-x_{ex})\right]\right]\cdot(2-s)\right] \div 80$$
$$f_{S,AS} = \frac{1}{3}\cdot s\cdot\left[24 + \left(23+\frac{1}{3}\right)\cdot(1-x_{ex})\right] \div 80$$

Eq. (A8)

where $x_{ex}$ is the same as above (degree of water exchange during lipid synthesis, and *s* is the degree of H contribution to lipids from substrate, $H_2$. Similar to the heterotrophic case above, the same four scenarios with end-member values of $x_{ex}$ and *s* are applied to the autotrophic case. Here, the three reduction steps involved in autotrophic Ac-CoA synthesis (Steps 1, 4 and 5; Fig. 2B) are the reverse of oxidative Ac-CoA degradation during heterotrophy (Steps 4, 5 and 8; Fig. 2A). The first reduction step (Step 1, Fig. 2B) involves $Fd_{red}$, and the next two reduction steps (Steps 4 and 5, Fig. 2B) involve $F_{420}H_2$, where H in both of these electron donors are incorporated directly via water ($f_{W,cofa-AS}$, Fig. 3B). For the autotrophic case, we assign the maximal substrate contribution (*s* = 100 %) when the $H_2$-dependent isoenzyme, Hmd, catalyzes the methenyl-$H_4$MPT reduction (Step 4, Fig. 2B) under high partial pressure of $H_2$ (see 4.1.2). In this case, one of the three methyl-H in Ac-CoA are incorporated via $H_2$ ($f_{S,AS}$, Fig. 3B).

### Culturing conditions for *Acidianus* sp. and *Metallosphaera sedula*

Batch cultures of *Acidianus* sp. DS80 and *Metallosphaera sedula* DSM 5348T were grown for additional archaeal lipid $\varepsilon_{L/W}$ measurements. *Acidianus* sp. DS80 was grown with $H_2$ as the electron donor as previously described. Briefly, axenic cultures of *Acidianus* sp. DS80 were grown on a defined mineral medium (Boyd et al., 2007) containing Wolfe's vitamin and SL-10 trace metals solutions. For the $H_2/Fe^{3+}/CO_2$ condition ('hf' in Figure 6), ferric iron ($Fe^{3+}$) was added in the form of ferric sulfate solution to a final concentration of 7 mM. For the $H_2/S^0/CO_2$ condition ('hs' in Figure 6), elemental sulfur ($S^0$) was sterilized by baking at 100 °C for 24 hours and added to the medium at a concentration of 5.0 g/L. For both conditions, medium was purged with $N_2$ gas followed by replacement of headspace with $H_2/CO_2$ (80:20, v/v). All cultures were incubated statically at 80 °C and an initial pH of 3.0, which remained within 0.1 units throughout the experiment. Axenic cultures of *M. sedula* DSM 5348T were grown on Brock basal salts medium containing Allen's trace element solution (Brock et al., 1972). The headspace of the culture bottles were flushed with $N_2$, then $CO_2$, $H_2$ and $O_2$ were added to the final pressure of 2 atm and final mixing ratios of 1–10 %, 30% and 5%, respectively. The pH of the medium was adjusted to 2.0. All cultures were incubated at 75 °C and 150 rpm.

### Full list of references for Figure 7

**(A)** $\varepsilon_{L/W}$ values observed in acetogenic and isoprenoid lipids from the domains Archaea and Eukarya. Different classes of compounds are shown on the horizontal axis). **For acetogenic pathway:** n-alkanol data for *Cryptomeria japonica* (Chikaraishi et al., 2004); *n*-alkane data for *Zostera marina* and *Spartina alterniflora* (Sessions et al., 1999); *n*-alkyl (*n*-alkanoic acids and fatty acids) data for *Alexandrium fundyense*, *Isochrysis galbana*, *Ascophyllum* sp., *Fucus veiculosis*, *Botryooccus braunii*, *Eudorina unicocca*, *Volvox aureus*, *Cucumis sativus*, *Emiliania huxleyi*, and *C. japonica* (Sessions et al., 1999; Chikaraishi et al., 2004, 2009; Zhang and Sachs, 2007; Sachs et al., 2016); alkadiene data for *A. fundyense*, *I. galbana*, and *B. braunii* (Sessions et al., 1999; Zhang and Sachs, 2007). **For the MVA pathway:** $C_{15}$ and $C_{30}$ isoprenoids (sterol, squalene, triterpenoids, and sesquiterpenes) for *A. fundyense*, *I. galbana*, *Ascophyllum* sp., *F. vesculosis*, *Z. marina*, *S. alterniflora*, *C. sativus*, *E. huxleyi*, and *C. japonica* (Sessions et al., 1999; Chikaraishi et al., 2004, 2009; Sachs et al., 2016); P+BP (phytanes and biphytanes) for *Nitrosopumilus maritimus*, *Sulfolobus acidocaldarius*, *Sulfolobus* sp., *Metallosphaera sedula*, *Acidianus* sp. DS80, *Methanosarcina barkeri*, and *Archaeoglobus fulgidus* (Kaneko et al., 2011; Wu et al., 2020; Leavitt, Kopf et al., 2023; this study). **For the MEP pathway:** $C_{20}$ isoprenoids (diterpenes, sandaracopimarinol, ferruginol, diterpenol, and sugio) for *C. japonica* (Chikaraishi et al., 2004); phytol+ptd (phytol and phytadiene) for *A. fundyense*, *S. alterniflora*, *B. braunii*, *Volvox aureus*, *C. sativus*, *E. huxleyi*, and *C. japonica* (Sessions et al., 1999; Chikaraishi et al., 2004, 2009; Zhang and Sachs, 2007; Sachs et al., 2016); and btc (botryococcene) for *B. braunii* (Zhang and Sachs, 2007). **(B)** $\varepsilon_{L/W}$ values observed in fatty acids from anaerobic and/or autotrophic Bacteria and isoprenoid lipids from Archaea. Different organisms are shown on the horizontal axis. **For Bacteria:** *E. coli* (Ec; Osburn et al., 2016), *Paracoccus denitrificans* (Pd; Osburn et al., 2016), SRB (*Desulfobacterium autotrophicum*, *Desulfococcus multivorans*, *Desulfobacter hydrogenophilus*, *Desulfovibrio alaskensis*, *Desulfolobus propionicus*; Campbell et al., 2009; Dawson et al., 2015; Osburn et al., 2016; Leavitt et al., 2016), and *Cupriavidus* sp. (Zhang et al., 2009). **For Archaea:** *Acidianus* sp. DS80 (Ac; this study), *Sulfolobus* sp. (Su; Kaneko et al., 2011), *Archaeoglobus fulgidus* (Af; this study), *Metallosphaera sedula* (Ms; this study), *Nitrosopumilus maritimus* (Nm; Leavitt, Kopf et al., 2023), and *Methanosarcina barkeri* (Mb; Wu et al., 2020).

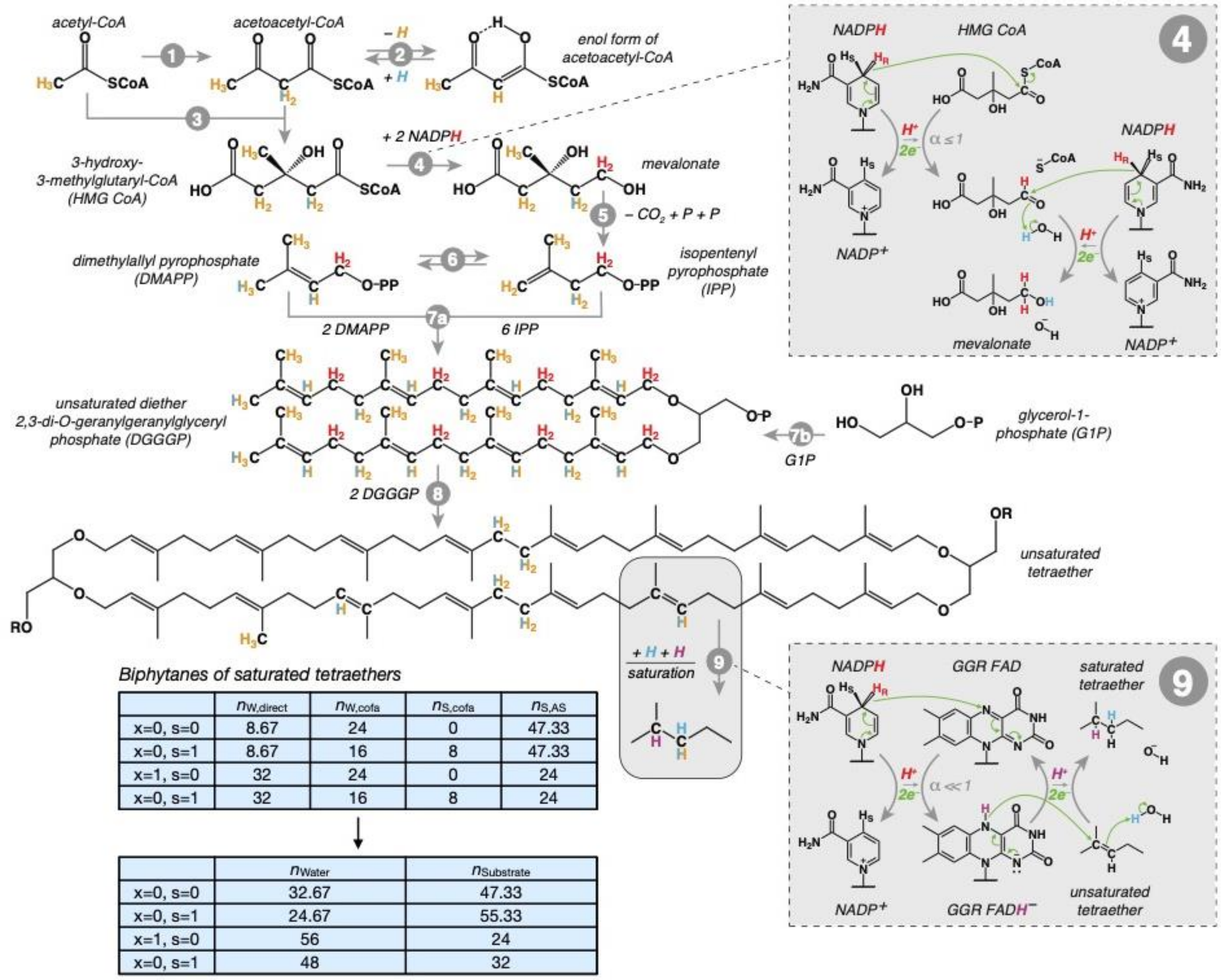


| | $n_{W,direct}$ | $n_{W,cofa}$ | $n_{S,cofa}$ | $n_{S,AS}$ |
|---|---|---|---|---|
| x=0, s=0 | 8.67 | 24 | 0 | 47.33 |
| x=0, s=1 | 8.67 | 16 | 8 | 47.33 |
| x=1, s=0 | 32 | 24 | 0 | 24 |
| x=0, s=1 | 32 | 16 | 8 | 24 |

| | $n_{Water}$ | $n_{Substrate}$ |
|---|---|---|
| x=0, s=0 | 32.67 | 47.33 |
| x=0, s=1 | 24.67 | 55.33 |
| x=1, s=0 | 56 | 24 |
| x=0, s=1 | 48 | 32 |

**Figure S1.** Overview of hydrogen sources for tetraether biosynthesis in *Archaeoglobus fulgidus* (figure modified from Leavitt, Kopf et al. (2023). Hydrogen atoms are color-coded according to their H sources: H from acetyl-CoA (orange); H from NADPH (red); H from water $H^+$ (blue). Mixed potential sources of H from acetyl-CoA and water are shown half orange/half blue. The summary boxes in blue indicates the number of biphytane H that could originate from water without going through substrate ($n_{Water}$) and the number of biphytane H via substrates ($n_{Substrate}$) during tetraether biosynthesis. These values depend on the extent of re-equilibration during isomerization (Steps 2 and 6) and contribution from organic substrate (lactate) during heterotrophy or $H_2$ during autotrophy. See Table 3 and Appendix A for details. The insets for steps 4 and 9 show the different hydride transfers. The GGR (step 9) is proposed to be highly fractionating. See the main text for further discussion. Biosynthetic steps are indicated in grey. 1: acetyl-CoA acetyl transferase; 2: tautomerization of acetoacetyl-CoA (can exchange the H at the $C_2$ position); 3: HMG-CoA synthase; 4: HMG-CoA reductase; 5: several alternative pathways from mevalonate to IPP (no H differences); 6: IPP isomerase (can exchange the H at the $C_4$ position); 7a: geranylgeranyl pyrophosphate synthase; 7b: geranylgeranyl glyceryl phosphate synthase; 8: tetraether synthase (Tes); 9: geranylgeranyl reductase (GGR).

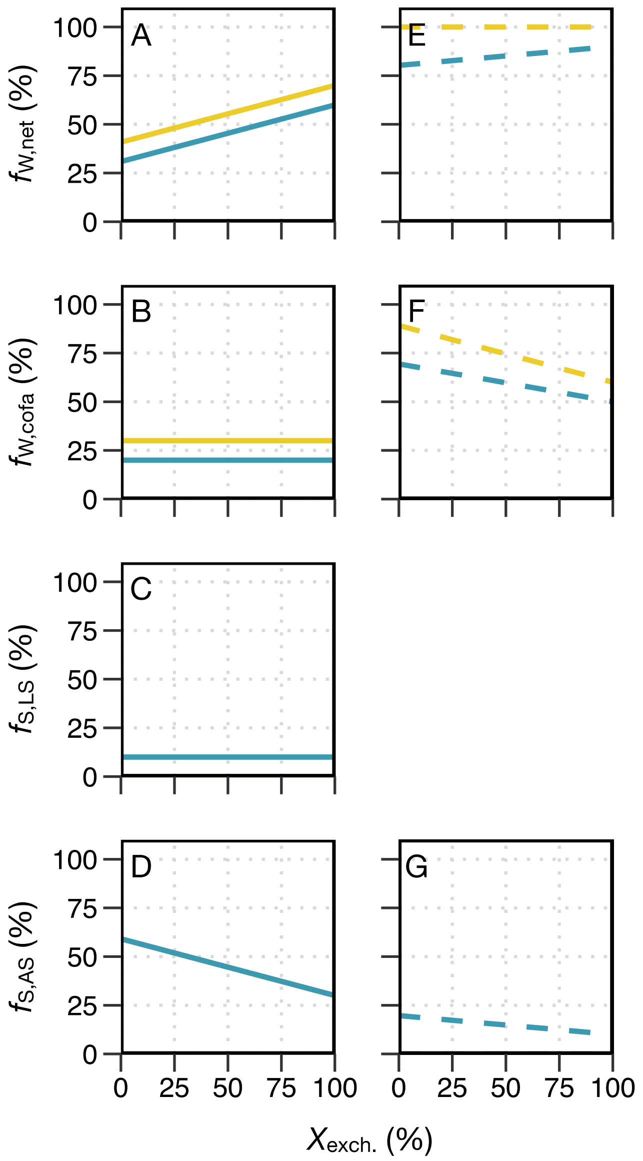


**Figure S2.** Isotope flux-balance model results for *Archaeoglobus fulgidus*. For all plots, the x-axis represents $X_{exch}$ or the degree of water exchange during lipid synthesis, which depends on the extent of re-equilibration during isomerization steps (Steps 2 and 6; Appendix A). Each column represents the metabolic mode: heterotrophy (either on thiosulfate or sulfate and lactate) (A–D); or autotrophy on thiosulfate and $H_2/CO_2$ (E–G). Each row represents one of the fluxes depicted in Fig. 3 in the main text: $f_{W,net}$ = total flux that directly or indirectly (e.g., via cofactors) reflects water isotope composition ($f_{W,net} = f_{W,direct} + f_{W,Cofa\text{-}LS}$ for heterotrophy; $f_{W,net} = f_{W,direct} + f_{W,Cofa\text{-}LS} + f_{W,Cofa\text{-}AS}$ for autotrophy) (A, E); $f_{W,cofa}$ = total flux of H that is incorporated into lipids via reducing cofactors (NADPH and GGR for heterotrophy; NADPH, GGR, Fd, and $F_{420}H_2$ for autotrophy) (B, F); $f_{S,LS}$ = total flux of H that is incorporated into lipids via organic substrate (lactate) during lipid synthesis (only applicable to heterotrophy) (C); and $f_{S,AS}$ = total flux of H that is incorporated into lipids via substrates (lactate for heterotrophy; $H_2$ for autotrophy) during acetyl-CoA synthesis (D, G). See Appendix A for details.
